# Structural basis of a regulatory switch in mammalian complex I

**DOI:** 10.1101/2023.12.14.571638

**Authors:** Daniel N. Grba, John J. Wright, Zhan Yin, William Fisher, Judy Hirst

**Affiliations:** Medical Research Council Mitochondrial Biology Unit, University of Cambridge, Keith Peters Building, Cambridge Biomedical Campus, Hills Road, Cambridge CB2 0XY, UK; Department of Biochemistry, University of Cambridge, Tennis Court Road, Cambridge, CB2 1GA, UK

## Abstract

Respiratory complex I powers oxidative phosphorylation in mammalian mitochondria by using the reducing potential of NADH to reduce ubiquinone-10 and drive protons across the inner mitochondrial membrane. High-resolution cryoEM structures have provided a molecular framework for complex I catalysis, but controversies about how to assign functional properties to the states identified in single-particle analyses are preventing progress on its energy-converting mechanism. Here, we combine precise biochemical definition with high-resolution cryoEM structures in the phospholipid bilayer of coupled vesicles and show that the closed and open states observed in mammalian complex I preparations are components of the deactive transition that occurs during ischaemia. Populations of the cryoEM open state and biochemical deactive state match exactly. Deactivation switches the enzyme off, converting the closed state that is capable of rapid, reversible catalysis into an open, dormant state that is unable to start up in reverse. The deactive state is switched back on by slow priming reactions with NADH and ubiquinone-10. Thus, by developing a versatile membrane system to unite structure and function, we define the role of large-scale conformational transitions in complex I and establish a new gold standard for structure-based investigations of catalysis by energy-coupled proteins.

---

Single-particle electron cryomicroscopy (cryoEM) has revolutionised our ability to determine high-resolution structures of large and complicated biological molecular machines, such as the mitochondrial energy-transducing complexes of oxidative phosphorylatio^1,2^. The capability of cryoEM to isolate multiple distinct states from heterogeneous mixtures lends new power to investigations of conformation-based mechanisms, but demands combination with biochemical and biophysical data to assign and position structurally defined states onto catalytic cycles. Respiratory complex I (NADH:ubiquinone oxidoreductase)^3–5^, one of the largest membrane-bound enzymes in the mammalian cell, powers ATP synthesis in mitochondria by using the reducing potential of NADH to drive protons across the inner membrane. CryoEM has delivered high-resolution structures of this L-shaped, 45-subunit complex and its homologues from multiple species, but structures observed in different states have been subjected to a wide variety of interpretations^6–19^. Furthermore, the diverse mechanistic proposals they have inspired are compromised because the structures have been determined in detergent micelles, and thus lack clear relationships with biochemical and biophysical data determined in physiologically relevant membrane environments that, unlike detergent micelles, support energy-conserving catalysis with native long-chain ubiquinones and transmembrane proton-pumping reactions. To solve the mechanism of complex I catalysis requires integration of structural and functional information into a single coherent and robust picture.

Here, we present high-resolution structures of mammalian (*Bos taurus*, bovine) complex I embedded in phospholipid vesicle membranes containing ubiquinone-10 (Fig. 1). In these complex I-containing proteoliposomes (CI-PLs) the enzyme can be switched between different states, and activated by addition of a partner ubiquinol-10 oxidase (the alternative oxidase, AOX) to perform rapid and reversible catalysis^20^ with rates corresponding to the maximum rates of turnover observed in native membranes (Fig. 2a)^21^. By adding different partner enzymes, CI-PLs can be modified further to catalyse NADH-driven ATP synthesis, or NAD^+^ reduction by reverse electron transfer (RET), both mediated by a substantial proton-motive force^22,23^. Here, we apply cryoEM to CI-PLs to define the structures of the ‘active’ and ‘deactive’ resting states of mammalian complex I that underpin its role in ischaemia–reperfusion injury^24–27^, establishing a new quality threshold for structural studies of complex I catalysis. Our structural assignments depend on the innovative combination of advanced biochemical, biophysical and cryoEM strategies (Extended Data Fig. 1): the preparation and reconstitution of highly active CI into CI-PLs suitable for cryoEM, a comprehensive biochemical definition of the enzyme states present, and an advanced application of single-particle cryoEM methods.

**Fig. 1.**
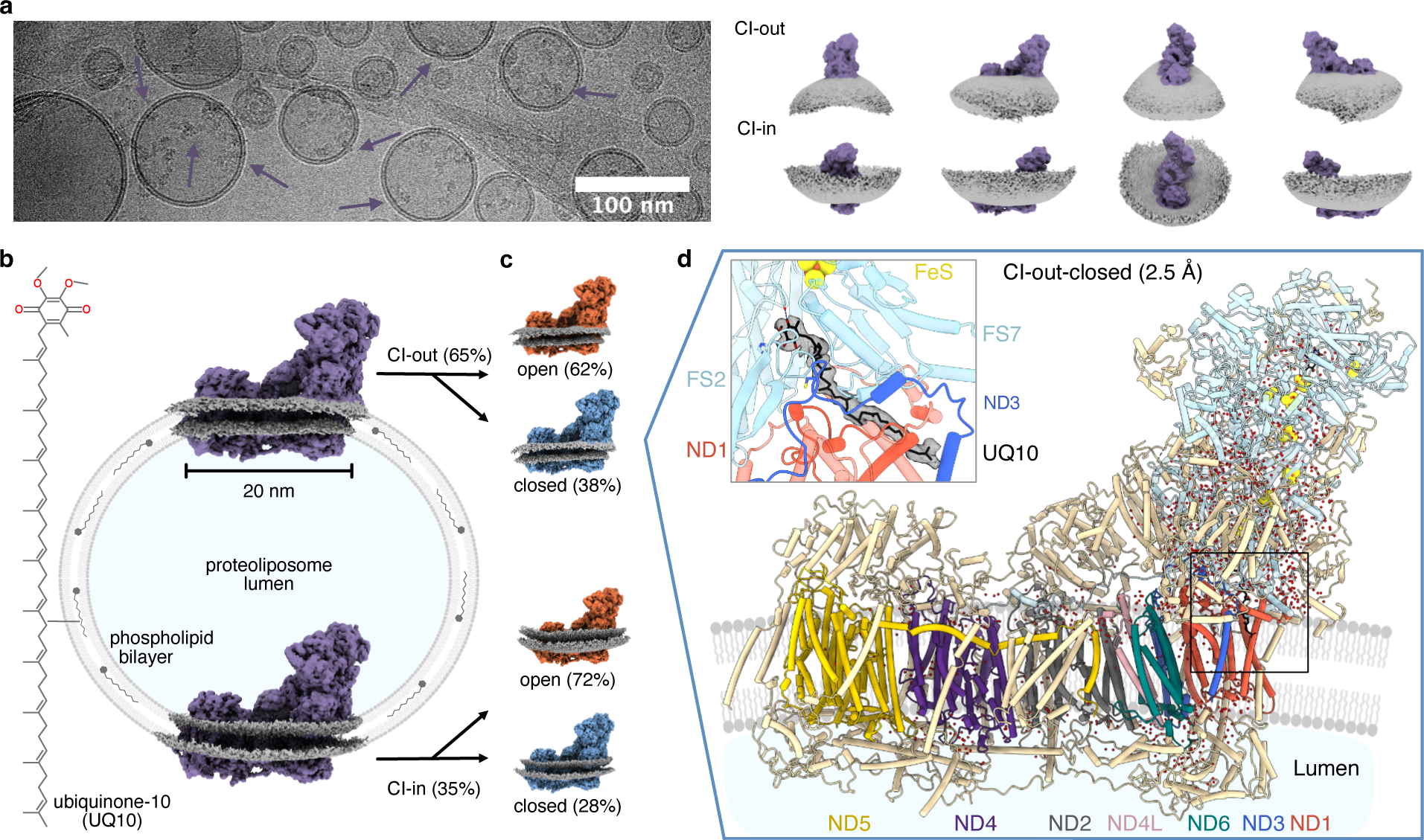
Generation of high-resolution structures for complex I in proteoliposomes. **a)** An example micrograph (CI in CI-PLs marked with arrows) and 3D volumes for CI embedded in CI-PLs facing outwards (CI-out) and inwards (CI-in). **b)** Visualisation of CI reconstituted into CI-PLs containing ubiquinone-10. The cryoEM density for the lipid bilayer (grey) around the enzyme (purple) was clipped and extrapolated to display a typical 50 nm-diameter liposome. **c)** CI-out and CI-in were classified into open (orange) and closed (blue) states and processed to generate high-resolution maps. **d)** The model for CI-out-closed (2.5 Å resolution) is shown with water molecules (spheres) and ubiquinone-10 in the substrate-binding site (inset). Conserved core subunits are in light blue (hydrophilic domain) or labelled in colour (membrane domain) and supernumerary subunits are in wheat.

**Fig. 2.**
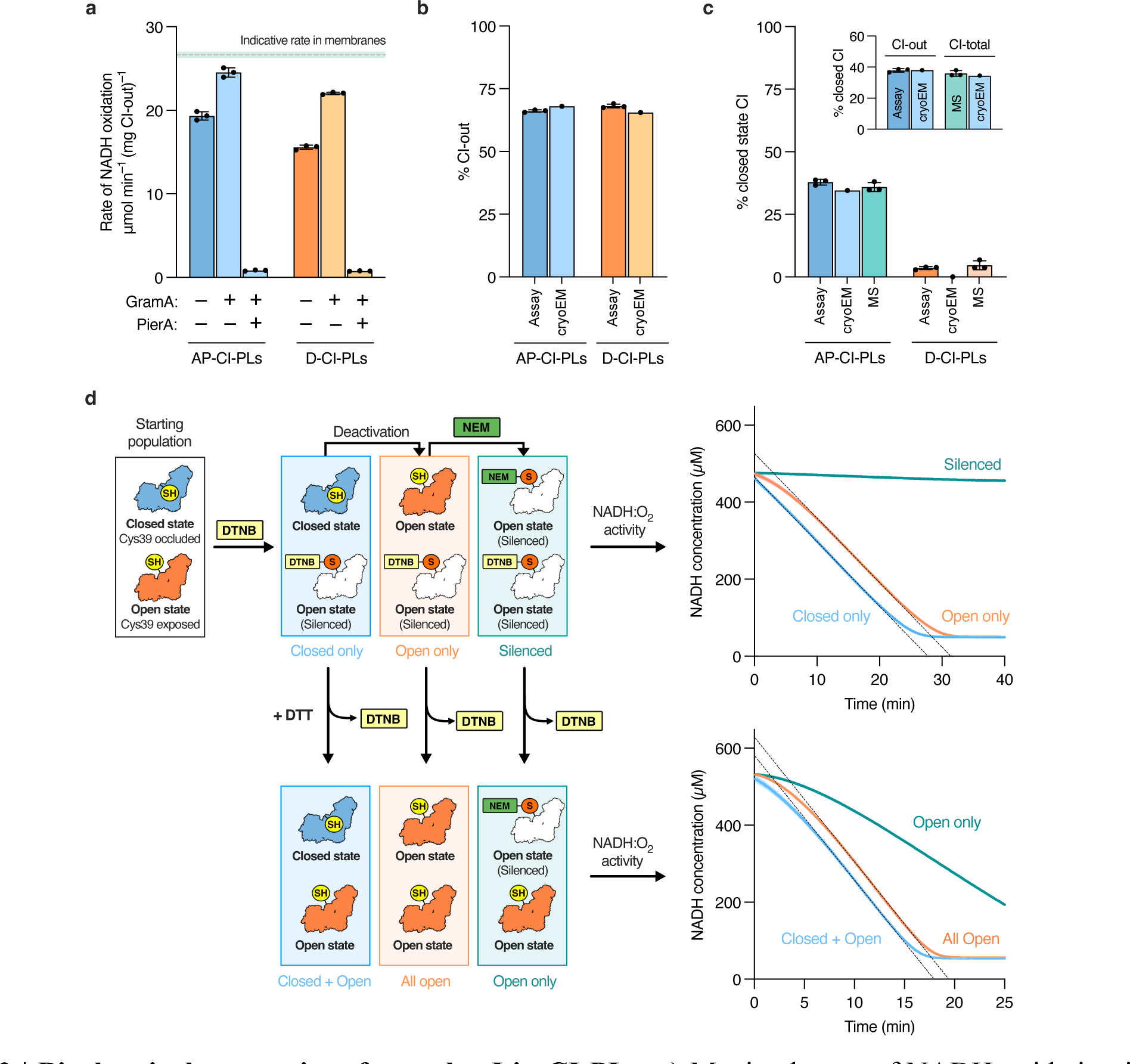
Biochemical properties of complex I in CI-PLs. **a)** Maximal rates of NADH oxidation in as-prepared (AP) and deactivated (D) cryoEM samples of CI-PLs measured with the ubiquinol oxidase AOX in the assay buffer (*n* = 3). Gramicidin A (GramA, to dissipate Δp, 0.5 µg mL^−1^) and piericidin A (PierA, to inhibit CI, 1 µM) were added as indicated. An indicative value for CI in mitochondrial membranes is provided for comparison, calculated on the basis that 10% of the protein present is CI^21^. **b)** Proportion of outward facing complex I (CI-out) in cryoEM samples determined using the NADH:APAD^+^ oxidoreduction assay (n = 3) and from the cryoEM particle distribution. **c)** The proportion of closed-state complex I in AP-CI-PLs and D-CI-PLs determined using the sensitivity of the rate of NADH:O_2_ oxidoreduction to NEM (assay, *n* = 3), by cryoEM particle classification (cryoEM), and by mass spectrometry following isotopic IAM labelling (MS, *n* = 3). MS data are from three independent CI-PL reconstitutions. The inset figure presents the values from the cryoEM data for AP-CI-PLs to match the CI-out and CI-total populations determined by biochemical assays and MS, respectively. **d)** The lag phase observed in complex I activity assays is due to the presence of open states (*n* = 3). A mixture of closed and open states in an as-prepared sample of CI-PLs was (i) treated with DTNB (only the closed population catalyses, linear catalysis), (ii) deactivated (the previously closed population opens, and catalyses with a lag phase), then (iii) labelled with NEM (both populations are inactive). Each sample was then treated with DTT to remove the DTNB. In (i) both states catalyse and the mixture displays a lag phase; in (ii) the whole population is open and the lag phase is increased; in (iii) only the open population catalyses, with a lag phase. See Extended Data Fig. 5g for a matching experiment using only IAM.

## Closed and open states in the membrane

CryoEM images of membrane-bound CI in the CI-PLs, flash frozen onto graphene oxide grids, were collected and separated into distinct populations (Fig. 1). The first classification separated CI molecules embedded with their large hydrophilic domain, which extends into the mitochondrial matrix in vivo, facing either out of the PL lumen (CI-out) or into the PL lumen (CI-in) (Fig. 1a-b). The proportion of CI-out is ∼65%, determined by both cryoEM classification encompassing the surrounding bilayer and by biochemical assays that rely on accessibility of the NADH-binding site (CI-out only, versus CI-total when the membrane is permeabilised) (Fig. 2b). In both orientations, density for the surrounding bilayer is clearly visible, curving away from the protein membrane domain (Fig. 1a), and allowing visualisation of the complex within a typical 50 nm-diameter proteoliposome (Fig. 1b). The second classification separated both CI-out and CI-in into two major conformational states, which we refer to as the ‘closed’ and ‘open’ states^6^ (Fig. 1c). Further classification procedures indicated the closed conformation is structurally homogenous, but separated the open state into two similar but distinct states that we refer to as open1 and open2. The three states (closed, open1 and open2) are related by rotation of the hydrophilic domain and ‘heel’ relative to the rest of the membrane domain (Extended Data Fig. 2a–c), which appears visually as an opening and closing of the angle between the domains, changing the secondary domain interface between NDUFA10 (membrane domain) and NDUFA5 (hydrophilic domain) substantially (Extended Data Fig. 2d)^6,28,29^. The highest resolution map (CI-out-closed) refined to 2.5 Å, and near-complete protein models were built to all six maps, together with a substantial number of ordered phospholipids and water molecules (Fig. 1d) (Supplementary Tables S1-2, Figs. S1–3). The maps and models show how amphipathic helices at the base of the hydrophilic domain that anchor the complex to the membrane interface influence the membrane topology, and how the packing of transmembrane helices (TMHs) in the membrane domain adjust the complex to the outward or inward curvature (Extended Data Fig. 3). Otherwise, the maps and models for outward- and inward-facing states are essentially identical. In the first part of our analysis, open1 and open2 (which are shown below to be indistinguishable biochemically) will be treated as a single ‘open’ state.

The closed and open structures observed here correspond to the closed and open structures of bovine CI determined previously in *n*-dodecyl β-D-maltoside (DDM) detergent micelles and in phospholipid nanodiscs prepared from the DDM-purified enzyme^9,28^ (Extended Data Fig. 4). However, the so-called ‘slack’ structure observed previously in both detergent and nanodiscs, which exhibits substantial disorder and disruption in key regions including subunit NDUFA11 and the C-terminus of ND5, could not be detected in CI-PLs. Having previously observed DDM instead of phospholipids bound behind the ND5 C-terminal transverse helix^16^, plus a DDM molecule inserted into the narrow ubiquinone-binding channel in the open state of bovine complex I in nanodiscs^9^, we have here switched the detergent used for purification to lauryl maltose neopentyl glycol (LMNG), which is equivalent to two DDM molecules joined together and is expected to be too bulky to enter the channel^7^. As the slack state was present in a matching analysis of CI-PLs formed from DDM-purified CI (DDM-CI-PLs, Supplementary Table S3, Figs. S4–6), and it was the only state observed in a catalytically inactive preparation from macaque heart^30^, we conclude it is an artefact promoted by detergent-induced displacement of specific bound phospholipids and/or detergent entering the ubiquinone-binding site. The importance of considering detergent effects was also highlighted in cryoEM analyses of complex I from the yeast *Chaetomium thermophilum* and the bacteria *Escherichia coli*, where purification in DDM or LMNG altered the distributions of states observed^12,13^.

## Assigning functions to structural states

Two competing models currently assign the closed and open states observed by cryoEM in preparations of mammalian CI to different frameworks. The first postulates they are both resting states^28,29,31^ (states that form post-catalysis in the absence of substrates). The second postulates they are both catalytic intermediates^6,12^ (representing CI molecules that halted at different points on the catalytic cycle, when substrate depletion or changing conditions stopped turnover). Further progress on understanding the mechanism of CI catalysis requires clarification of these models. The biochemical characteristics of the resting states of mammalian CI have long been established^24,26,32^. The active (A) ready-to-go resting state can immediately begin forward (NADH:ubiquinone oxidoreduction) or reverse (RET) catalysis upon introduction of substrates so it displays linear (constant-rate) steady-state NADH oxidation catalysis. The A state is also insensitive to thiol-reactive reagents that derivatise Cys39^ND3^ on the TMH1–2^ND3^ loop adjacent to the ubiquinone-binding site^33,34^. In contrast, the deactive (D) dormant resting state requires substrate-induced reactivation to catalyse in the forward direction, exhibiting an initial catalytic ‘lag phase’ during its reactivation, and it is unable to start catalysing in reverse^23,24^, a feature that is considered protective against oxidative damage^25,27^. In the D state Cys39^ND3^ is sensitive to derivatisation, which prevents reactivation by locking the enzyme in a catalytically inactive state^33,34^. The A state converts slowly and spontaneously to the D state at elevated temperature in the absence of substrates for turnover^24^.

In our preparation of CI-PLs (‘as-prepared’ CI-PLs), cryoEM classification determined that 35% of the molecules are closed (65% are open), with a slightly higher proportion closed in the outward-facing population (Figs. 1c and 2c). The cryoEM values were compared to data from three biochemical analyses for the A and D states. First, we used two methods that rely on Cys39^ND3^ only being accessible in the D state^23,34^. In a standard approach, we treated the CI-PLs with a thiol-reactive reagent and used activity assays (which report only on CI-out, as only its NADH-binding site is accessible) to determine the proportion of enzyme that was irreversibly inactivated. Three thiol-reactive reagents, *N*-ethyl maleimide (NEM), iodoacetamide (IAM), and 5,5′-dithiobis-(2-nitrobenzoic acid) (DTNB), all gave similar results (Extended Data Fig. 5a). We also used a sequence of isotopic labelling and mass spectrometry to measure total (CI-out and CI-in combined) Cys39^ND3^ labelling directly^35^ (Extended Data Fig. 6). In both cases, the biochemical proportions of A and D are equal to the cryoEM proportions of the closed and open states (Fig. 2c), consistent with expectations from structure models that show Cys39^ND3^ is occluded in the closed state, but exposed and solvent accessible in open states (Fig. 3a)^6,28,29,31,36^. Second, we tested for the catalytic lag phase that is a well-established biochemical feature of the D state, using alkaline conditions to prolong reactivation^24,26,33^. We used DTNB to derivatise Cys39^ND3^ reversibly (the DTNB label can be removed by treatment with the disulphide-reducing agent dithiothreitol (DTT), Extended Data Fig. 5b^37^) and a standard thermal deactivation of CI-PLs (37 °C for 20 min, Extended Data Fig. 5c) to convert all the enzyme to the D state^23,31^. The data show conclusively that the catalytic lag phase is only observed when substrates are supplied to the (underivatised) open enzyme, and that catalysis is linear with no lag phase when only the closed enzyme is able to catalyse (Fig. 2d). Deactivated CI-PLs (D-CI-PLs) show all the expected biochemical characteristics of the D state, including full recovery of their catalytic activity following reactivation and derivatisation of Cys39^ND3^ in activity assays and mass spectrometry analyses (Fig. 2a–c and Extended Data Figs. 5b–d), and in cryoEM analyses all the CI molecules were in the open state (Supplementary Table S4, Figs. S7–10). Furthermore, the open1 and open2 structures determined in D-CI-PLs were indistinguishable from those determined in the as-prepared CI-PLs (Extended Data Fig. 7) so the deactive enzyme generated by deactivation is identical to the deactive enzyme present (at lower levels) in as-prepared samples. Our results confirm unambiguously that the cryoEM closed state is the ready-to-go ‘active’ resting state and the cryoEM open states are the dormant, deactive resting states; the resting states are formed post-catalysis in the absence of substrates and are not catalytic intermediates.

**Fig. 3.**
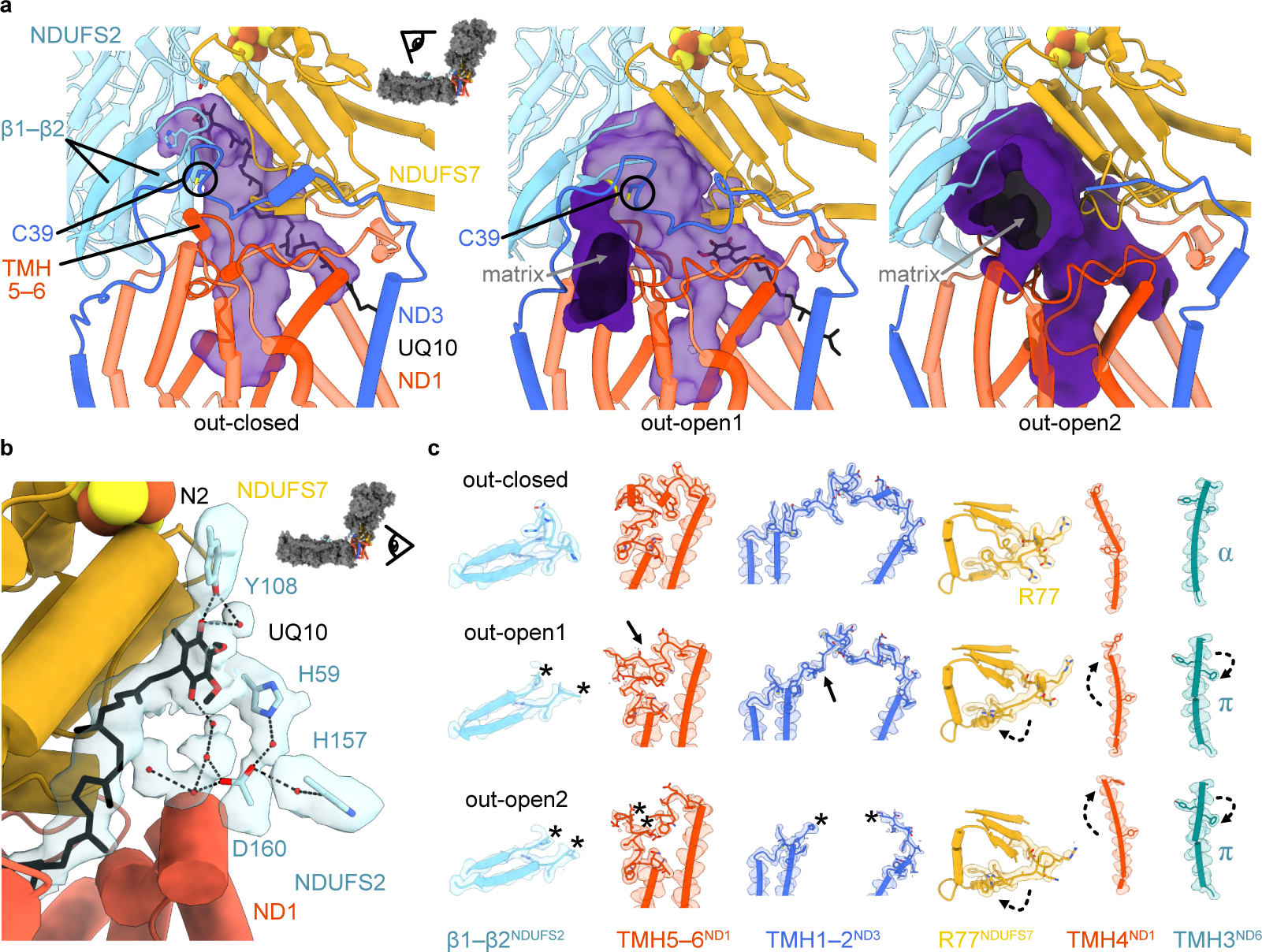
Local features of complex I structures in CI-PLs. **a**) The ubiquinone-10 (UQ10)-binding channel in the closed, open1 and open2 states. Closed: the UQ10-binding channel (purple, semi-transparent surface) is sealed from the matrix, UQ10 is fully inserted, and C39^ND3^ is buried from the matrix. Open1: the UQ10-binding channel is sealed from the matrix, UQ10 is partly inserted, and C39^ND3^ is exposed to the matrix (purple, solid surface with yellow indicating the contribution from Cys39^ND3^). Open2: the UQ10-binding channel is exposed to the matrix (purple, solid surface), no bound UQ10 is present, and C39^ND3^ is exposed to the matrix. **b)** The bound UQ10 in the CI-out-closed state reveals a two hydrogen-bonding networks of water molecules (red spheres) that complete its connection to the proposed ligands (His59^NDUFS2^ and Tyr108^NDUFS2^) for protonation upon reduction by the terminal iron–sulfur cluster N2. The semi-transparent surface shows the cryoEM density (2 Å distance, threshold value 0.03). **c**) The local elements that correlate with closed and open structural states: β1–β2^NDUFS2^ loop (ordered in closed, progressively less ordered in open1 and open2); TMH5–6^ND1^ loop (ordered in closed and open1, minor disorder in open2), TMH2–3^ND3^ loop (ordered in closed, ordered and shifted in open1, disordered in open2), R77^NDUFS7^, TMH3,^ND6^ α helix/π bulge and TMH4^ND1^. CryoEM density is shown (at thresholds 0.1, 0.1, 0.09, 0.1, 0.1, 0.09, left to right) with residues of interest shown as sticks. Solid and dashed arrows indicate localized disorder and movement, respectively, relative to the closed state. Asterisks indicate peptides not observed due to disordering. See also Extended Data Table 1.

## The ubiquinone-10 binding site

The CI-in/out-closed structures exhibit well-ordered density for ubiquinone-10, inserted fully into its active site with one of the ubiquinone carbonyls directly hydrogen bonded to Tyr108^NDUFS2^, a proposed ligand and proton donor for ubiquinone reduction^38,39^ (Fig. 3b). Focussed classifications on the binding site were unable to separate any substates, suggesting the site is saturated by a single ubiquinone-10 binding pose, consistent with its free equilibration with the ∼10 mM ubiquinone-10 in the PL bilayer (relative to *K*_M_ values of 0.5−1.5 mM for catalysing CI-PLs^22,23,40^). Earlier deep-bound ubiquinone-10 structures have either been in a pre-reactive conformation lacking direct interaction with the Tyr108^NDUFS2^ ligand^9^ or without sufficient resolution to model water molecules^11^. Here, water molecules are observed around the ligated ubiquinone headgroup, in a single hydrogen-bonding network that indirectly connects both proposed proton donors (His59^NDUFS2^ and Tyr108^NDUFS2^) to both ubiquinone carbonyls (Fig. 3b). Otherwise, the local, detailed features (Fig. 3c, top and Extended Data Table 1) and discrete networks of residues and water molecules within the E-channel and hydrophilic axis (Extended Data Fig. 8) match previous closed-state mammalian structures^9,16^. The tightly structured closed ubiquinone-binding site at the hydrophilic–membrane domain interface likely determines this consistency.

In as-prepared and deactivated CI-PLs the open state was resolved into two similar states, open1 and open2, that are distinguished by differences in the status of loops at the ubiquinone-binding site and domain interface, but otherwise conform to all the established structural hallmarks of open, deactive states (Fig. 3c and Extended Data Table 1)^41^. In open1, the TMH1–2^ND3^ loop is ordered in a slightly altered conformation to in the closed state, accommodating and also restricting global opening. At the deepest site in the ubiquinone-binding channel, the β1–β2^NDUFS2^ loop is disordered and the trigonal interaction^14^ between Cys39^ND3^, His55^NDUFS2^ and Tyr127^ND1^ has been lost. Importantly, while Cys39^ND3^ is solvent accessible and available for derivatization by thiol-labelling reagents in the open1 state, the ubiquinone-binding channel is sealed from the matrix (Fig. 3a). The open1 structure thereby combines an open global conformation with a closed ubiquinone-binding site. In the lower section of the channel the open1 TMH5– 6^ND1^ loop is ordered in its closed-state conformation, and ubiquinone-10 is partially inserted into the open1 channel (its headgroup is adjacent to Glu202^ND1^), whereas in the open2 state both the TMH1–2^ND3^ and β1–β2^NDUFS2^ loops are disordered (Cys39^ND3^ is again accessible) and the ubiquinone-binding channel is open to the matrix (Fig. 3a). Although only a poorly defined density is observed, perhaps suggestive of weakly bound ubiquinone-10, the TMH5–6^ND1^ loop remains in its closed-state conformation, with a minor region of increased disorder at residues 208–209, alongside increased local disorder in the TMH3–4^ND6^ loop (residues 79–85) and NDUFA9 C-terminal domain (residues 325–332), relative to in the open1 substate. The ordered status of the TMH5–6^ND1^ loop in CI-PLs, which is conserved in our open2-like structure in DDM-CI-PLs, is in striking contrast to the extended disorder observed previously in open states in detergent micelles and nanodiscs^6,9,31,41^ (Extended Data Table 1). The results suggest that the local conformational changes induced by DDM are reversible upon DDM removal and enzyme reconstitution into CI-PLs. Finally, the presence of two distinct states within the open population in CI-PLs raises the question of whether both states are canonically deactive: both are sensitive to thiol-derivatizing reagents, but because we cannot control their formation independently, we cannot confirm whether both states or just one cause the open-state-dependent lag phase in catalytic activity assays. Therefore, we exploited the fact that deactive CI is unable to catalyse RET and assessed the ability of our CI-PLs to catalyse RET in a co-reconstituted system^23^ that uses F_1_F_O_ ATP synthase to create a proton-motive force (Extended Data Fig. 9). As-prepared CI-ATPase-PLs exhibited 35% of the maximum RET rate (achieved following a pre- reaction activation step), which matches the proportion found to be in the closed state using the NEM assay, showing that none of the open enzyme is able to catalyse RET. Furthermore, no RET could be observed for deactivated CI-ATPase-PLs, which contain both the open1 and open2 states only. Therefore, open1 and open2 are both deactive states, consistent with the ND6 π-bulge disconnecting proton transfer from the ubiquinone-10 binding site^6,9^ and blocking RET^42^.

## Structural basis for deactivation

Spontaneous but slow transition of the closed resting state into deactive open states indicates deactivation is an exergonic process with a high activation energy, but how the structural changes that occur define the energy landscape for the transition and confer these properties is not understood. Here, we consider how the membrane environment influences the stabilities of the states and the transition between them. Comparing the membrane-bound closed and open states, the hydrophilic domain rotates about an axis projected on subunit ND1 through the membrane domain (Fig. 4a and Extended Data Fig. 2a–b), with ND1 and surrounding supernumerary subunits as a pivot point. TMH2–8^ND1^, supported underneath the hydrophilic domain by NDUFA8, NDUFA1 and NDUFA13, bend as the enzyme closes, or extend as it opens (Fig. 4a–b). The long, curved NDUFA13 helix bends or extends alongside, whereas NDUFA8 and NDUFA1 are fixed and appear to brace the pivoting of NDUFA13 at its helix-breaking Pro-72 residue. We propose TMH2–8^ND1^ extending and straightening provide the primary driving force for deactivation, with their bending and deformation back into closed, catalytically active states demanding energisation. Opposing straightening of these loaded helical ‘springs’, interaction of the amphipathic NDUFA9 C-terminal domain with the lipid bilayer creates a counterbalance (Fig. 4c). In the closed state, an abundance of well-ordered NDUFA9-associated lipid molecules embed and anchor it to the bilayer whereas, in open states, rotation of the hydrophilic domain lifts NDUFA9 away from the bilayer and far fewer lipids are observed. On the opposite side of the domain, amphipathic helices on NDUFS8 and NDUFA12 are pushed further into the bilayer in open states, creating an intrinsic stabilising ‘buoyancy’. Additional protein interactions stabilizing the closed state^43^ include the greater buried interface between NDUFA5 and NDUFA10 (Extended Data Fig. 2d) and the N-terminus of NDUFS2 that runs from the hydrophilic domain across the matrix surface of the membrane domain, where it is ordered in the closed state but disrupted and disordered in open states.

**Fig. 4.**
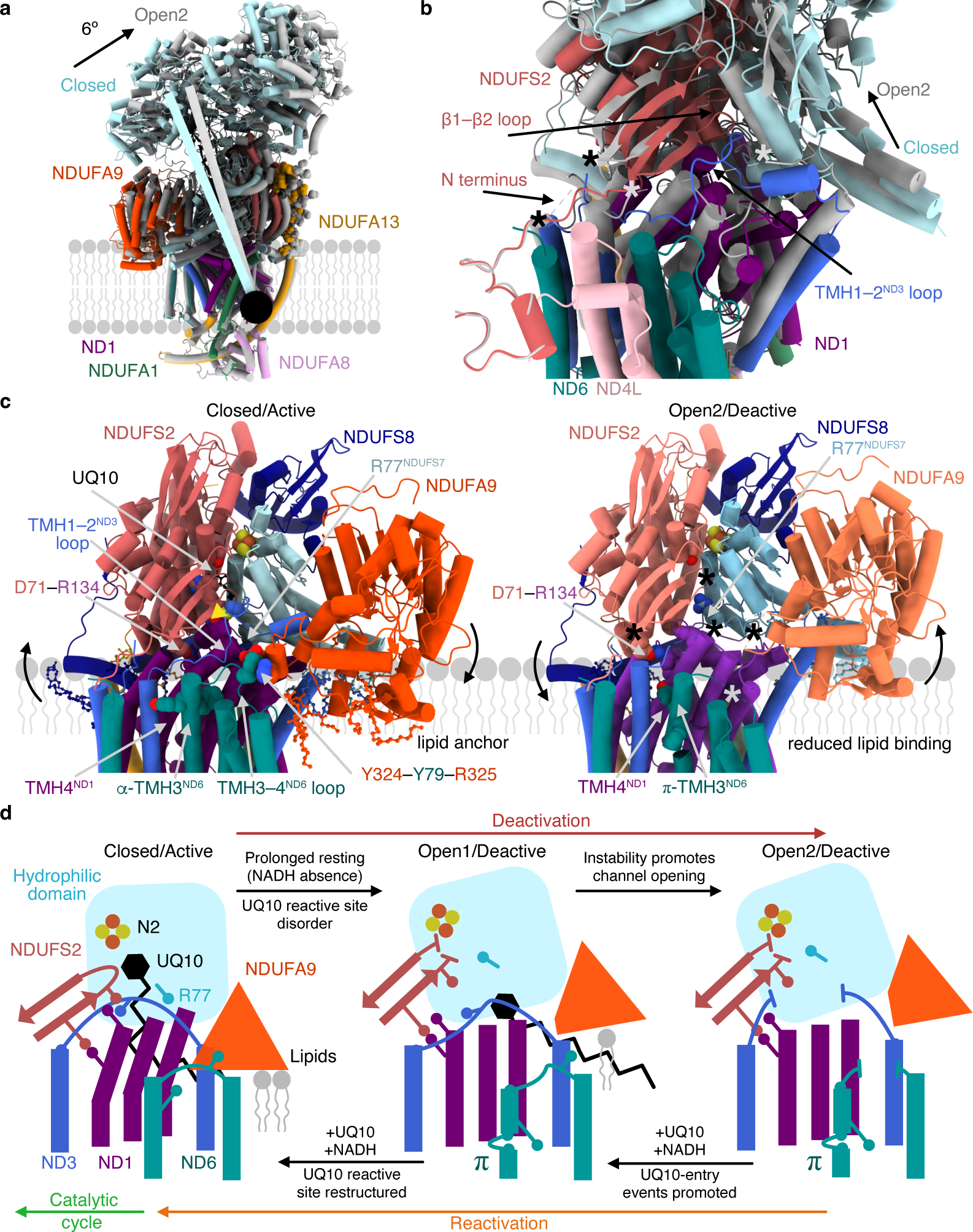
Structural transition between the active/closed and deactive/open resting states. **a)** Global comparison of the closed/active (colour) and open2/deactive (grey) structures superpositioned on their membrane domains showing relevant supernumerary subunits from behind the hydrophilic–membrane domain interface. The axis of rotation is marked as a black circle, with the rotation (6°) planes shown. The N-terminus of NDUFA13 extends up the hydrophilic domain and is shown as spheres. **b)** The same alignment as in **a** viewed from the front and focused on the hydrophilic-membrane domain interface to show the movement of the ND1 TMHs relative to ND6 and the rotation of the hydrophilic domain. Asterisks indicate disorder in the open2/deactive model, where coordinated colours link the missing residues. **c)** The same superpositioned models as in **b** with labelled subunits of interest only and the closed/active and open2/deactive states shown side by side. Lipids and ubiquinone-10 (UQ10) are shown in ball and stick representation; iron–sulfur cluster N2 and key residues are shown as spheres. The residues of the trigonal junction are indicated by the yellow triangle. Asterisks indicate regions of loop disorder. **d)** Schematic overview of the structural elements involved in the deactivation and reactivation transitions between the closed/active and open2/deactive states corresponding to panel **c**. Circles on sticks indicate stabilizing residue interactions, such as the trigonal junction of H55^NDUFS2^, Y127^ND1^ and C39^ND3^; R77^NDUFS7^, the ion pair R134^ND1^–D71^NDUFS2^; and the packing interactions Y324^NDUFA9^–Y79^ND6^– R325^NDUFA9^ and F68^TMH3-ND6^ with ND1.

Ubiquinone-10 fully inserted into its binding channel in the closed state stabilises the β1–β2^NDUFS2^ loop that carries the essential His59, as well as the closed position of Arg77^NDUFS7^ (since it clashes sterically with its open position). Upon deactivation Arg77^NDUFS7^ moves into the channel, blocking ubiquinone-10, bound part-way into the channel in the open1 state, from transiting deeper in. Excluding substrate from the deep-bound site allows the β1–β2^NDUFS2^ loop to escape its closed conformation, encouraging His55^NDUFS2^ to break its trigonal interaction with Cys39^ND3^ and TMH4-Tyr127^ND1^ as the latter straightens (Fig. 4c). Asp71 on the β1–β2^NDUFS^^2^ loop remains tethered to TMH4-Arg134^ND^^1^ by an ion-pair interaction as the straightening ND1-TMHs drive rotation of the hydrophilic domain, generating the matrix-exposed Cys39^ND3^ cavity observed in open1. Further destabilisation of the TMH1–2^ND3^ loop, which crosses the rotation interface, occurs upon conversion to open2 (Fig. 3a and Fig. 4b–d). Both straightening of the ND1-TMHs and rotation-induced displacement of NDUFA9 destroy the closed-state packing site for TMH3-Phe68^ND6^, and destabilise interactions between TMH3–4-loop-Tyr79^ND6^ and Tyr324– Arg325^NDUFA9^, promoting the helical rotation of TMH3^ND6^ that generates the hallmark π-bulge of the open states (Figs. 3c and 4c–d). The π-bulge (with unfavourable dihedral angles, decreased hydrogen-bond interactions and decreased internal packing) has an intrinsic driving force to return to an α-helical conformation, aiding reactivation. Prolonged absence of fully-bound ubiquinone-10 from the closed state thereby promotes the coordinated, exergonic opening transition of a network of interacting structural elements, which are otherwise finely counterbalanced to maintain the system in the metastable closed states in which catalysis occurs. Returning deactive open states to catalysis requires energy input from NADH-driven ubiquinone-10 reduction, to refold and reform local elements and drive domain rotation, to regenerate the strained, loaded helical springs in ND1 and re-anchor NDUFA9 to the lipid bilayer, re-poising the system for energy transduction (Fig. 4d).

## Discussion

CI-PLs enable high-resolution cryoEM analyses of fully functional complex I, in its native membrane environment, to be combined with coherent and precise biochemical characterisation using its native ubiquinone-10 substrate. Here, we used CI-PLs to show that the structurally defined closed and open resting states of mammalian complex I are the biochemically defined active (ready-to-go) and deactive (dormant) resting states. They are well-behaved, distinct states that can be prepared and interconverted reversibly with full conservation of catalytic activity, consistent with their physiological relevance.

The closed state in CI-PLs contains ubiquinone-10 fully loaded into the catalytic site, reflecting its ready-to-go nature. The open1 and open2 substates are both deactive off-cycle resting states that are dormant and must be re-activated to return to catalysis, with their off-cycle status confirmed by their inability to initiate reverse catalysis. Our robust and coherent model for the role of the open states is not consistent with the idea, based on the presence of open states in cryoEM analyses of as-prepared samples of ovine complex I in detergent micelles, that they are catalytic intermediates^6^. Furthermore, in contrast to our consignment of the slack state of bovine complex I, containing extensive disorder in the C-terminal section of ND5 and adjacent subunit NDUFA11, to a detergent-induced artefact, equivalent substates of ovine complex I in detergent micelles were included in the heterogeneous open states proposed to represent the catalytic intermediates^6^. The inability of these artefactual states to reactivate^30^ suggests why the relative populations of the closed and open ovine states did not respond to the addition of substrates, and the persistence of slowly reactivating open states also does not suggest they are catalytic intermediates^6^. Finally, our data confirm that purposeful deactivation of as-prepared samples of complex I in membranes merely converts the closed enzyme present into the same open enzyme already there at lower levels. Our data are not consistent with the alternate ‘deactive’ structure created by heat-treating the ovine enzyme in detergent micelles^6^, which was presented without substantive biochemical data to confirm its identity or ability to return to catalysis, and which exhibited considerable structural disorder and alteration.

The proposed role of supernumerary subunit NDUFA9 in stabilising the closed state of the mammalian complex through extensive bilayer-phospholipid binding (Fig. 4), suggests how delipidation by membrane-disrupting detergents may drive it into open states. Similarly, amphipathic helices in the hydrophilic-domain core subunits likely stabilise complexes, such as *E. coli* complex I, which lack NDUFA9 and do not display the biochemical characteristics of mammalian-type active/deactive transitions. Artefactual delipidation may explain the dominance of pronounced open states in cryoEM analyses of DDM-purified *E. coli* complex I^8^, and the appearance of the “open-ready” state, which most resembles our open1 state conformation, when the milder LMNG detergent was used^12^. For *E.coli* complex I, the necessity of closing the open states for catalysis, supported by appearance of the closed state on addition of substrates to preparations that were otherwise all open^12^, may explain the delay in starting proton pumping observed after the onset of redox catalysis^44^. The possibility of detergent-induced structural artefacts compromises confidence in mechanistic, biochemical and physiological interpretations from detergent-bound structures and underlines the need to transfer future cryoEM studies of complex I catalysis into membrane environments.

Although our data do not support open states participating directly in catalysis, the coordinated global changes of the deactive transition may provide clues to the catalytic mechanism. The same local elements that change their conformation together during deactivation may change individually during catalysis, but in a local and constrained fashion^45^, without breaking open the ubiquinone-binding site (and thereby incurring the energy costs of de-solvation, refolding and resealing) or globally opening the enzyme. Here, our open1 state, which is a globally open state with a sealed ubiquinone-binding site, demonstrates that even these two effects are not incontrovertibly linked together. Open1 is ‘inbetween’ the closed and open2 states, and we suggest it forms first upon deactivation, and is the predominant deactive state formed in vivo (with open2 promoted by in vitro conditions that encourage solvent influx to the ubiquinone-binding site, destabilising any ubiquinone-10 present). Open1 is likely also an intermediate in reactivation of open2, formed when substrate entry to the channel begins to order local elements, encouraging substrate transit further into the channel before energy input from electron transfer from NADH drives the site to reform and globally close. In our model, ubiquinone-10 binding is essential for refolding disordered loops and reforming the site during reactivation, whereas substrate exchange for catalysis occurs within closed-only states. Our model is consistent with observations of quinone bound partly or fully inserted into the closed channel in mycobacterial complex I^18^, and with molecular dynamics simulations that have illustrated its transit through the closed channel on physiologically relevant timescales^46,47^. We thus conclude it is not necessary to form open2-like states during catalysis, to open the ubiquinone-binding site to the matrix to allow waters to flow in and out to accommodate or replace the substrate, as was proposed previously to justify their assignment to the catalytic cycle^6,12^. The conflicting interpretations of structural data discussed here underline the need for robust functional analyses to be employed alongside structural analyses in future mechanistic studies, as enabled by our CI-PLs.

Our study sets a new gold standard for cryoEM studies on respiratory complex I, and highlights new possibilities for mechanistically challenging systems that require membrane environments for their function and/or suffer from poor distribution and orientation biases in their detergent-solubilised forms. Structural work on proteoliposomes for other target proteins has already been demonstrated in seminal work by Yao and coworkers^48^, who reported the first medium-resolution (3.8-Å) cryoEM structure of the AcrB multidrug resistant transporter, embedded in liposomes on graphene grids. Structures of several ion channels have now also been determined and separated into different states in liposomes^49^, and a recent 2.7-Å resolution structure of a high conductance Ca^2+^-activated K^+^ channel has recently been determined in native membrane vesicles^50^. Here, we demonstrate high resolution structures and discrete states for a non-symmetrical protein, using grid preparation methods that require only standard equipment. The native environment and ubiquinone-10 substrate provided in CI-PLs, combined with the exquisite biochemical control that the system allows, has finally clarified the enigma of the closed and open states, and allowed a sophisticated structural evaluation of the resting states of mammalian complex I and their functional relevance. The CI-PLs system will now provide unprecedented opportunities for cryoEM studies of the catalytic intermediates that are generated under catalysing and energy-transducing conditions.

## Methods

### Purification of *Bos taurus* complex I in LMNG

Mitochondrial membranes were prepared from bovine heart tissue as described previously^51^. Complex I was purified using a method adapted from the established procedure for purification in *n*-dodecyl-β-D-maltopyranoside (DDM)^40^. Membranes (175 mg protein) were treated with one cOmplete protease inhibitor tablet (Roche) then solubilised on ice at 4.5 mg mL^−1^ by the stepwise addition of lauryl maltose neopentyl glycol (LMNG, Anatrace) from a 5% stock solution to a final concentration of 1.5%, followed by a further 20 min incubation, and centrifugation (8,500 × g, 15 min, 4 °C). The supernatant was filtered through a 0.22 µm syringe-tip filter then loaded onto three 5 mL Q-Sepharose HP columns (Cytiva) connected in series and pre-equilibrated with Buffer A (20 mM Tris-HCl (pH 7.5 @ RT), 2 mM EDTA, 10% (v/v) ethylene glycol, 0.15% (w/v) LMNG, 0.02% asolectin (total soy lipid extract, Avanti Polar Lipids), and 0.02% 3-[(3-cholamidopropyl)-dimethylammonio]-1-propanesulfonate (CHAPS)). The columns were washed with 23% Buffer B (Buffer A + 1 M NaCl) then Buffer B was increased to 38% to elute complex I. Complex I-containing fractions were collected, pooled and concentrated using an Amicon Ultra-15 (100 MWCO) then injected onto a Superose 6 Increase column (Cytiva) pre-equilibrated with Buffer C (20 mM Tris-HCl (pH 7.5 @RT), 150 mM NaCl, 10% (v/v) glycerol, 0.0375% LMNG). Complex I-containing fractions were concentrated as before and flash frozen in liquid N_2_. The complex I concentration (according to the bicinchoninic acid (BCA) assay) and the NADH:APAD^+^ and NADH:DQ oxidoreductase rates were determined.

### Purification of AOX

AOX from *Trypanosoma brucei brucei* was expressed recombinantly in *E. coli* and purified from membranes by solubilisation with octyl-glucoside and Twin-Strep tag affinity chromatography as described previously^40^.

### Preparation of CI-PLs

Complex I proteoliposomes (CI-PLs) for cryoEM and biochemical analyses were prepared using a method adapted from a previously published procedure^23^. 1,2-dioleoyl-*sn*-glycero-3-phosphocholine (DOPC), 1,2-dioleoyl-*sn*-glycero-3-phosphoethanolamine (DOPE), and 18:1 cardiolipin (CDL) were mixed at an 8:1:1 weight ratio as a 25 mg mL^−1^ chloroform stock solution. 10 mg of lipids were transferred to a glass tube and supplemented with 100 nmol Q_10_, then the chloroform was removed using a gentle stream of N_2_. The resulting lipid film was transferred to a desiccator and stored under vacuum for at least 1 hr. Lipids were rehydrated with frequent vortexing over 30 min following addition of 1 mL of Proteoliposome Buffer (10 mM MOPS-KOH (pH 7.5, 20 °C), 50 mM KCl). Small unilamellar vesicles (< 50 nm) were formed by sonication on ice using a Q700 probe sonicator (qSonica) equipped with a 1.6 mm microtip (60% amplitude, 2.5 min, cycle of 15 s on, 30 s off) then partially solubilised by addition of 0.5% (w/v) sodium cholate and incubated on ice for 10 min. Complex I was added at a 50:1 (w/w) lipid to protein ratio and the mixture incubated on ice for a further 20 min before the detergent was removed using a PD10 desalting column (Cytiva) pre-equilibrated with Proteoliposome Buffer. CI-PLs were collected by ultracentrifugation (210,000 × g, 1 hr, 4 °C) and resuspended in the same buffer. The resulting sample was centrifuged using a benchtop centrifuge at maximum speed (16,000 × g, 10 min, 4 °C) to remove any larger particles, and kept on ice or at 4 °C before use. All the data reported for CI-PLs are from a single preparation of complex I to ensure that data from the different experiments can be compared. Biochemical and mass spectrometry analyses on multiple samples of CI-PLs, from different reconstitutions of the same complex I, were confirmed to be consistent.

### Complex I activity assays

All activity assays were performed spectrophotometrically using a SpectraMax plus 348 or SpectraMax ABS Plus 96-well plate reader (Molecular Devices). Unless otherwise stated, assays were conducted at 32 °C. Complex I concentration in CI-PLs was determined using the NADH:APAD^+^ oxidoreduction reaction (ε_400−450_ = 3.16 mM^−1^ cm^−1^): the standard rate was determined for LMNG-solubilised complex I (0.5 µg mL^−1^) in Proteoliposome Buffer with 100 μM NADH, 500 μM APAD^+^, 500 nM piericidin A and 0.2% (w/v) DDM^22,40,52^, and compared to the rate from CI-PLs under the same conditions but without the DDM, giving the concentration of CI-out. The total concentration of CI, and proportion of CI-out, were determined by comparing the NADH:APAD^+^ rates in the presence and absence of 15 µg mL^−1^ alamethicin (AlaM) to permeabilise the proteoliposomes to the substrates. NADH:O_2_ oxidoreduction rates were measured as the rates of NADH oxidation (ε_340−380_ = 4.81 mM^−1^ cm^−1^) in the presence of excess AOX to continually re-oxidise the ubiquinone-10^20^. CI-PLs (0.5 µg mL^−1^ CI-out) were diluted into Proteoliposome Buffer and treated with 5–10 µg mL^−1^ AOX, then catalysis was initiated by addition of 200 µM NADH.

The uncoupled rate (Δp = 0) was calculated following addition of 0.5 µg mL^−1^ gramicidin A (GramA). Proton pumping was assessed using the fluorescence of 9-amino-6-chloro-2-methoxyacridine (ACMA) (λ_ex_ = 410 nm, λ_em_ = 480 nm) in a RF-5301PC spectrofluorometer (Shimadzu) at 32 °C. CI-PLs (1.0 µg mL^−1^ CI-out) were incubated in Proteoliposome Buffer with 10 µg mL^−1^ AOX, 0.5 μM ACMA and 0.1 μM valinomycin and 500 µM NADH added to initiate catalysis. When required, 1 µM piericidin A was added to inhibit complex I, or 15 µg mL^−1^ alamethicin for uncoupling. Reverse electron transfer assays were performed with proteoliposomes co-reconstituted with both CI and *E. coli* F_1_F_O_ ATP synthase (3 ATPase per CI) (Extended Data Fig. 9), supplemented with glycerol 3-phosphate dehydrogenase (GlpD, *E. coli*) as described previously^23^ but with minor modifications to the activation procedure. To activate complex I, CI-PLs (10 µg mL^−1^ CI-out) were incubated at 32 °C with 60 µM NADH for 2 min, then 60 µM decylubiquinone (DQ) was added. After a further 1 min incubation, during which the NADH and DQ were consumed, RET was initiated by addition of ATP, *rac-*glycerol 3-phosphate and NAD^+^ as described previously^23^. Assay data are reported ± SD with individual data points shown.

### Derivatisation of Cys39 to determine A/D ratios

To assess the proportion of A-state complex I relative to the total amount of (A+D) enzyme present, CI-PLs were diluted to 200 µg mL^−1^ and treated with either 1 mM *N-*ethylmaleimide (NEM, from a 100 mM stock in 25% DMSO), 20 mM iodoacetamide (IAM, from a 200 or 500 mM stock in Proteoliposome Buffer) or 0.2 mM 5,5′-dithiobis-(2-nitrobenzoic acid) (DTNB, from a 10 mM stock in DMSO). Samples were incubated on ice for 20 min (NEM), 30 min (IAM) or 1 hour (DTNB) to ensure complete labelling. Where indicated, complex I was deactivated by incubating the CI-PLs at 37 °C for 20 min. NADH:O_2_ oxidoreductase activities were determined as above, following dilution of the sample into Proteoliposome Buffer and addition of AOX and NADH. To determine the A/D states in bovine mitochondrial membranes, 2 mg mL^−1^ membranes were treated with 1 mM NEM, then the activity determined in 10 mM Tris-HCl (pH 7.5), 250 mM sucrose using 5 µg mL^−1^ membranes, 3 µM cytochrome c (equine heart) and 15 µg mL^−1^ AlaM, with catalysis initiated by 200 µM NADH. Membranes were deactivated using the same procedure as for CI-PLs.

For differential isotopic labelling for mass spectrometry studies, freshly prepared samples of CI-PLs (typically 250–350 µg mL^−1^ CI-out) were split into three: an ‘as-prepared’ (AP-CI) sample, a deactivated (D-CI) sample that was incubated at 37 °C for 20 min, and a vehicle control AP-CI sample that was treated identically to the AP-CI sample but without inclusion of the labels. The samples were incubated with 30 mM H-IAM (Label 1, L1) on ice for 30 min, then the H-IAM was then quenched by addition of 50 mM GSH for 10 min on ice, and the CI-PLs were collected by centrifugation (210,000 × g, 1 hr, 4 °C). The CI-PL pellets were each washed with 3 × 1 mL of Proteoliposome Buffer then resuspended in 100 µL of the same buffer. The NADH:O_2_ oxidoreductase activity of each sample was measured, and a portion of each sample was incubated with 20 mM L-IAM and its activity checked again to confirm completion of labelling. All the samples were then incubated at 37 °C for 20 min to deactivate them, and their activities checked to ensure that no denaturation had occurred and that no L1 had been carried over. They were then treated with 50 mM L-IAM (Label 2, L2) and incubated for 30 min on ice before the activities were recorded again. The samples were then divided into 20 µL aliquots, and each aliquot diluted with either 100 µL MS control buffer (20 mM Tris-HCl (pH 8.0), 200 mM NaCl, 100 mM GSH) for method 1, or 100 µL MS denaturation buffer (20 mM Tris-HCl (pH 8.0), 200 mM NaCl, 8 M urea, 50 mM L-IAM) for method 2, and stored at –20 °C. A schematic diagram of the labelling procedure is given in Extended Data Fig. 6a.

For trypsin cleavage, labelled samples (20 µL) were precipitated with 20 volumes of ice-cold ethanol overnight. The pellets were then digested with 20 µL of a 1:100 dilution of trypsin in 50 mM NH_4_HCO_3_. 2 µL of each digested sample was then added to 10 µL of 0.1% TFA, 3% CH_3_CN, and injected for LC/MS/MS analysis. The peptide mixtures were fractionated by liquid chromatography using a gradient of 5–40% CH_3_CN, 0.1% (v/v) TFA, at a flow rate of 300 nL min^−1^ over 84 min using an Acclaim PepMap nanoViper C18 reverse-phase column in an EASY-nLC system (Thermo Scientific). Fractions were transferred to a Q-Exactive Plus Orbitrap mass spectrometer (Thermo Scientific, UK) where data were acquired for a peptide mass range of 400 to 1600 m/z for precursor ions and the ten most abundant precursor ions were fragmented by high-energy collisional dissociation (HCD) in N_2_. For protein identification, the fragment patterns were compared to the UniProt database in Mascot. Relative quantitation was achieved by comparing the peak height of different extracted ion chromatograms (XICs) using Xcalibur.

To investigate catalytic lag phases, as-prepared CI-PLs (150 µL, 0.5 mg mL^−1^ CI-out) were incubated with 0.2 mM DTNB on ice for 1 h, then the sample was diluted with 1 mL Proteoliposome Buffer and centrifuged (210,000 × g, 1 h, 4 °C). The pellet was washed with 3 × 1 mL of Proteoliposome Buffer then the DTNB-labelled CI-PLs were resuspended in 100 µL of the same buffer. Half the labelled sample was deactivated (20 min incubation, 37 °C), and half the deactivated sample was treated with NEM (1 mM, 30 min incubation on ice). Assays to assess the lag phase were performed at room temperature in a buffer at elevated pH (10 mM Tris-HCl (pH 8.8), 50 mM KCl) to slow down reactivation and amplify the lag phase^26,33^. CI-PLs containing 2.5 µg mL^−1^ CI-out were combined with 10 µg mL^−1^ AOX and incubated at room temperature for 5 min prior to addition of NADH. Where indicated, 1 mM DTT was present to remove the DTNB label. For IAM labelled experiments, reconstituted CI-PLs were split into two, with one half of the sample incubated with L-IAM (30 mM, 30 min incubation on ice) and the other half of the sample untreated. In both cases, the lag phases were compared between as-prepared and deactivated CI-PL samples, with no DTT treatment.

### CryoEM grid preparation, screening and imaging

CryoEM grids [Cu grids (1.2/1.3 Quantifoil) for DDM datasets or UltrAuFoil® gold grids (0.6/1, Quantifoil) for LMNG datasets] were washed in chloroform and then ethanol, dried in air then glow discharged (20 mA, 90 s, 0.1 Bar). A solution of graphene oxide (GO, 2 mg mL^−1^, Sigma-Aldrich) was bath sonicated to evenly resuspend it, then 3 µL were applied to the top side of a grid and manually blotted. Grids were washed three times with water then dried at 100 °C for 30 min. Prior to sample addition, the GO-coated grids were glow discharged (10 mA, 15 s, 0.1 Bar). CI-PLs (2.5 µL per grid, 200 µg mL^−1^ CI-total) were applied to each grid in an FEI Vitrobot Mark IV (Thermo Fisher Scientific) at 4 °C and 95% relative humidity. The sample was incubated on the grids for 30 s then blotted for 2 s under a force setting of 0, and frozen by plunging into liquid ethane. Grids were screened at the cryoEM facility at the Department of Biochemistry, University of Cambridge, using either a Talos Arctica or a Titan Krios microscope (Thermo Fischer Scientific). Grids with good GO coverage, even ice and even proteoliposomes distribution were taken forward.

### As-prepared CI-PLs data collection

High-resolution images (27,014 micrographs) were recorded using a Titan Krios microscope at the cryoEM facility at the Department of Biochemistry, University of Cambridge. The instrument was operated at 300 kV using a Gatan K3 detector in super-resolution electron counting mode at a nominal sampling rate of 1.066 Å pix^−1^. Non-gain normalized images were acquired with AFIS and a post-column imaging energy filter (Gatan BioContinuum) set to a slit width of 20 eV at a dose rate of ∼17 electrons Å^−2^ s^−1^. The sample was exposed to a total dose of ∼45 electrons Å^−2^ in a 2.67 s exposure over 40 fractions. The C2 and objective lens apertures were set to 70 and 100 µm, respectively. Images were collected with a defocus range of −0.9 to −2.3 µm.

### DDM-CI-PLs data collection

High-resolution images (5,568 micrographs) were recorded using a Titan Krios microscope at the cryoEM facility at the Department of Biochemistry, University of Cambridge. The instrument was operated at 300 kV using a Gatan K3 detector in super-resolution electron counting mode at a nominal sampling rate of 1.43 Å pix^−1^. Non-gain normalized images were acquired with AFIS and a post-column imaging energy filter (Gatan BioContinuum) set to a slit width of 20 eV at a dose rate of ∼12 electrons Å^−2^ s^−1^. The sample was exposed to a total dose of ∼40 electrons Å^−2^ in a 3.40 s exposure over 40 fractions. The C2 and objective lens apertures were set to 70 and 100 µm, respectively. Images were collected with a defocus range of −1.0 to −3.0 µm.

### Deactivated CI-PLs data collection

High-resolution images (6,692 micrographs) were recorded using a Titan Krios microscope at the UK National electron Bio-Imaging Centre (eBIC), at the Diamond Light Source. The instrument was operated at 300 kV using a Gatan K3 detector in super-resolution electron counting mode at a nominal sampling rate of 1.072 Å pix^−1^. Non-gain normalized images were acquired with AFIS and a post-column imaging energy filter (Gatan BioContinuum) set to a slit width of 20 eV at a dose rate of ∼ 10.5 electrons Å^−2^ s^−1^. The sample was exposed to a total dose of ∼40 electrons Å^−2^ in a 3.8 s exposure over 40 individual movie frames. The C2 and objective lens apertures were set to 50 and 100 µm, respectively. Images were collected with a defocus range of −0.9 to −2.3 µm. An additional dataset (4,550 micrographs) was collected on the same grid with the same conditions except the stage was tilted by 20°, as suggested by cryoEF to improve orientation distribution^53^, and AFIS was disabled. The session reference is BI22238-45.

### As-prepared CI-PLs data processing

CryoEM data processing was carried out in RELION-4.0-beta^54^ and cryoSPARC v3.3.2^55^. Beam-induced motion in the gain-normalized movie frames was corrected using RELION’s implementation^56^ of MotionCor2^57^ and per-micrograph contrast transfer function (CTF) parameters determined from the non-dose-weighted motion-corrected movie sums with CTFFIND-4.1^58^. Initial particles (1,320,319) from on-the-fly processing using Warp^59^ were imported into cryoSPARC and a 200,000 subset subjected to multiple rounds of 2D classification to generate an initial subset of particles (16,358) to train the Topaz picker^60^ in RELION. This picker was then used on all motion-corrected, CTF-corrected micrographs and the above process was iterated until ∼6 million particles were picked. These particles were processed in cryoSPARC and RELION using a mixture of 2D and 3D classification and heterogenous and *ab initio* refinements facilitated here and throughout with the *csparc2star* python script of the UCSF pyem code repository^61^. A final set of 315,578 particles was then used to train Topaz before 5,688,318 particles (figure-of-merit value –4) were extracted (3.75 Å pix^−1^) and subjected to the final classification scheme (Supplementary Fig. S1). Here, the large number of particles were split into random groups and processed in parallel, before being combined at checkmark points. Heterogeneous refinement with six classes separated CI-in and CI-out using low-resolution membrane curvature features. Like classes were combined in homogeneous refinement before being randomly split again and applying iterative heterogenous refinements to remove junk particles and better sort the classes. The particles were re-extracted (1.066 Å pix^−1^, box size of 450 pixels) and randomly split within their respective CI-in/CI-out branches before protein-masked 3D classification (10 classes) and homogeneous refinement were run on each class. Masks were auto-generated in cryoSPARC. Classes were inspected for global conformation and grouped, with heterogenous jobs used to clean out junk particles. The particles were polished and CTF refined in RELION to facilitate subsequent classification in cryoSPARC using 3D classification and heterogeneous and homogeneous refinement. The classification resulted in two major protein states for each of CI-in and CI-out, open and closed, which were combined and re-extracted in RELION. Individual classes underwent particle polishing and CTF refinement prior to 3D classification in RELION over six classes, with angular sampling decreased gradually (7.5°/3.7°/1.8°) and local searches enabled, to help remove any remaining junk particles and verify the global classification from cryoSPARC through comparison of 3D refinement volumes of each class. We attempted a focus-revert-classify method (align on the hydrophilic arm prior to classification without alignment on the membrane arm^36^ to separate out the protein further); however, we found the result tended to identify different membrane curvatures resulting from different liposome sizes. Instead, we implemented a focus-subtract-classify-revert method. Briefly, particles were focus refined on the hydrophilic domain (to remove membrane arm-imposed limitation on alignment of the hydrophilic arm), revealing the conformational distribution at the membrane interface. The hydrophilic mask used was generated in RELION (with a binarization threshold of 0.02, an extension of three pixels and a soft cosine edge of 10 pixels) using a rigid-body fitted previous bovine complex I model (PDB 7QSL) and a volume generated using the *molmap* command in ChimeraX^62^, with the low-pass filter set to 15 Å. Particles were subtracted using a mask that covers the membrane– hydrophilic interface region, before being classified without alignments over eight classes using a regularization parameter (*T*) = 100. The mask of the region was generated as for the hydrophilic mask, except it was low-pass filtered to 10 Å, with a binarization threshold of 0.1 and an extension of five pixels. Output classes were reverted to their non-subtracted states and 3D refined before being combined (CI-in/out closed, open1 or open2) where appropriate for a final consensus 3D refinement. We found this method particularly effective for complex I; for example, it was able to separate out a sub-class of ∼10,000 particles from ∼335,000 particles that was overlooked by all other 3D classification methods performed above. Furthermore, it enabled us to separate the open state into discrete populations, termed open1 and open2, which differ in their protein loop ordering at the membrane–hydrophilic interface.

Global resolution was estimated in RELION from the Fourier Shell Correlation (FSC) between two independent, unfiltered half-maps, according to the FSC = 0.143 criterion^63^, and post-processed using an automatically estimated B factor: CI-out-closed = 2.5 Å, CI-out-open1 = 2.6 Å, CI-out-open2 = 2.6 Å, CI-in-closed = 2.7 Å, CI-in-open1 = 2.9 Å, CI-in-open2 = 2.6 Å. Local resolution was estimated in RELION. Maps were locally sharpened using MonoRes^64^ and LocalDeblur^65^ in the Scipion v1.2.1 package^66^ to aid in model building and model refinements. Map visualization was improved with *map_to_structure_factors* using Phenix v1.20.1-4487^67^. Directional resolution anisotropy was calculated using 3DFSC Program Suite v3.0^68^. Orientation distributions were plotted in Mollweide projections using Matplotlib and Python.

### DDM-CI-PLs data processing

CryoEM data processing was carried out in RELION-4.0-beta and cryoSPARC v3.3.2. Beam-induced motion in the gain-normalized movie frames was corrected using RELION’s implementation of MotionCor2 and per-micrograph contrast transfer function (CTF) parameters determined from the non-dose-weighted motion-corrected movie sums with CTFFIND-4.1. An initial manual picking was performed to train the Topaz picker in RELION and the picker was improved through the selection of good particles using 2D classification and heterogenous refinement, until a Topaz picker was trained on 410,047 particles. The picker was then used with a figure-of-merit value of −5 to select 2,313,863 particles for extraction at 4 Å pix^−1^. The particles were split into random subsets and processed in parallel, before being combined at checkpoints. Iterative heterogeneous refinement with six classes separated the CI-in and CI-out from junk classes via low-resolution membrane curvature features. Like classes were combined and homogenously refined before re-extraction (1.43 Å pix^−1^, box size of 360 pixels). A two-class 3D classification was performed in cryoSPARC with a mask around the membrane to better split CI-out and CI-in. Protein-masked 3D classification (six classes) was performed within the CI-out/CI-in branches. Masks were auto-generated in cryoSPARC. Classes were inspected for their global conformation and grouped, with heterogenous jobs used to clean out junk particles. A three-class 3D classification was then performed that provided three distinct classes: closed, open2 and slack. Particles were polished and CTF refined in RELION prior to 3D classification in RELION over six classes, with angular sampling decreased gradually (7.5°/3.7°/1.8°) and local searches enabled, to help remove any remaining junk particles and verify the global classification from cryoSPARC by comparing the 3D refinement volumes of each class. The resulting non-junk classes were refined and subjected to the focus-subtract-classify-revert classification scheme outlined above, split over four classes. The output classes were refined and grouped, resulting in six classes: CI-out/in-closed, CI-out/in-open2, CI-out/in-slack. Global resolution was estimated in RELION from the FSC between two independent, unfiltered half-maps, according to the FSC = 0.143 criterion, and post-processed using an automatically estimated B factor and to a calibrated pixel size of 1.35 Å pix^−1^, based on map–model correlation values with previous complex I models: CI-out-closed = 3.1 Å, CI-out-open2 = 3.1 Å, CI-out-slack = 3.6 Å, CI-in-closed = 3.1 Å, CI-in-open2 = 3.1 Å, CI-in-slack = 3.8 Å. Local resolution was estimated in RELION. Directional resolution anisotropy was calculated using 3DFSC Program Suite v3.0. Orientation distributions were plotted in Mollweide projections using Matplotlib and Python.

### Deactivated CI-PLs data processing

CryoEM data processing was carried out in RELION-4.0-beta and cryoSPARC v3.3.2 for both tilted and non-tilted datasets. In both cases, beam-induced motion in the gain-normalized movie frames was corrected using RELION’s implementation of MotionCor2 and per-micrograph contrast transfer function (CTF) parameters determined from the non-dose-weighted motion-corrected movie sums with CTFFIND-4.1. For both datasets, a previously trained Topaz picking model, using 801,932 particles from the as-prepared CI-PLs dataset, was implemented for particle picking in RELION. For the non-tilted dataset, a Topaz figure-of-merit value of 0 was used to select 878,425 particles for extraction at 2.86 Å pix^−1^. The particles were split into random subsets and processed in parallel, before being combined at checkpoints. Iterative heterogeneous refinement with six classes separated CI-in and CI-out via low-resolution membrane curvature features. Like classes were combined in homogeneous refinement, before being randomly split again and iterative heterogenous refinements were applied to remove junk particles and better sort the classes. Like classes were combined in homogeneous refinement before a final non-junk, six-class heterogeneous refinement was performed. Like classes were combined in homogeneous refinement and re-extracted (1.072 Å pix^−1^, box size of 480 pixels). Masks were auto-generated in cryoSPARC. No further processing in cryoSPARC was performed as no states other than the open state were detected in 3D classification or heterogeneous refinements (unlike in the as-prepared dataset). CI-in and CI-out particles were refined, polished and CTF refined in RELION prior to 3D classification in RELION over six classes, with angular sampling decreased gradually (7.5°/3.7°/1.8°) and local searches enabled, to help remove any remaining junk particles. Good classes were refined and subjected to a focus refinement and classification on the subtracted hydrophilic–membrane interface; however, this only removed minor junk classes. At this point the CI-out-open (87,008) and CI-in-open (40,990) particles were combined with equivalent particles from the tilted dataset.

For the 20°-tilted dataset, a Topaz figure-of-merit value of −2 was used to select 932,186 particles for extraction at 2.86 Å pix^−1^. The particles were split into random subsets and processed in parallel, before being combined at checkpoints. Iterative heterogeneous refinement with 12 classes separated the CI-in and CI-out via low-resolution membrane curvature features; the class number was increased as the low resolution of the tilted data made separating empty membrane/junk particles more difficult. Good protein classes were combined in homogeneous refinement before being randomly split again and iterative heterogenous refinements were applied to remove junk particles and better sort the classes. Good protein classes were combined in homogeneous refinement and re-extracted (1.072 Å pix^−1^, box size of 480 pixels) to improve resolution prior to a three-class heterogeneous refinement in which CI-out and CI-in were separated. They each underwent homogeneous refinement and a three-class heterogeneous refinement again before a final homogeneous refinement clean-up step. Masks were auto-generated in cryoSPARC. As in the non-tilted dataset, no further processing in cryoSPARC was performed as no states other than the open state were detected in 3D classification and heterogeneous refinements. CI-in and CI-out particles were refined, polished and CTF refined twice in RELION prior to 3D classification in RELION over six classes, with angular sampling decreased gradually (7.5°/3.7°/1.8°) and local searches enabled, to help remove any remaining junk particles. In each case, only a single good class was output, and then refined. At this point the CI-out-open (58,376) and CI-in-open (39,423) particles were combined with equivalent particles from the non-tilted dataset.

Prior to combining, the tilted and non-tilted particles were assessed for adjustments in orientation sampling (Supplementary Fig. S11). The tilt applied appeared to have reduced the orientation preference observed in the consensus refinement; however, this came at the cost of higher resolution. The high sphericity in the non-tilted dataset refinements suggests the orientation distribution does not suffer from a problematic lack of views. The combined CI-out-open (145,384) and CI-in-open (80,413) states underwent 3D refinement in RELION prior to 3D classification over six classes, with angular sampling decreased gradually (7.5°/3.7°/1.8°) and local searches enabled. The resulting non-junk classes [CI-out-open (142,957) and CI-in-open (76,4214)] were refined and subjected to the focus-subtract-classify-revert classification scheme outlined above, split over eight classes. The output classes were refined and grouped, with four classes resulting: CI-out/in-open1 and CI-out/in-open2. No closed states were observed. Global resolution was estimated in RELION from the FSC between two independent, unfiltered half-maps, according to the FSC = 0.143 criterion, and post-processed using an automatically estimated B factor: CI-out-open1 = 2.8 Å, CI-out-open2 = 2.6 Å, CI-in-open1 = 3.3 Å, CI-in-open2 = 2.9 Å. Local resolution was estimated in RELION. Maps were locally sharpened using MonoRes and LocalDeblur in the Scipion v1.2.1 package to aid in model building and model refinements. Map visualization was improved with *map_to_structure_factors* using Phenix v1.20.1-4487. Directional resolution anisotropy was calculated using 3DFSC Program Suite v3.0. Orientation distributions were plotted in Mollweide projections using Matplotlib and Python.

### As-prepared CI-PLs model building and refinement

Previous bovine complex I models were used as starting points for models for the closed (PDB 7QSK) and open (PDB 7QSN) states of complex I in the as-prepared CI-PLs^9^. They include the polymorphisms at NDUFA10-255 and NDUFS2-129 modelled as Lys and Arg^9,69–71^, respectively. Sequence numbering for subunits NDUFA6 and NDUFA13 were adjusted to account for N-terminal processing^69^. The cofactor in NDUFA10 was updated to Mg^2+^-bound 2’-deoxyguanosine 5’-triphosphate^72^. The models were rigid-body fit into respective consensus maps using ChimeraX v1.16.1^62^. Subunits were all-atom refined and per-chain refined using Coot v0.9.8.1^73^, guided by locally sharpened maps. ND5-locally refined maps, with a mask around respective CI-in/out-open/closed ND5 regions, were generated to improve the local resolution (Extended Data Fig. 3c) and used to build the variably curved ND5 regions with more confidence for each state, before reverting back to consensus maps. The ND5 region had relatively poor local resolution due to variation in liposome curvature. Previous lipid/detergent/ubiquinone-binding site molecules were removed from the starting PDB models, before lipids and substrate ubiquinone-10 molecules were built into the maps for each state. Added molecules were clipped where the cryoEM density was absent due to flexibility. Protein chains were inspected, and residues removed where cryoEM density was lacking, or added, including N-terminal acetylation, where densities not previously observed were present (see Supplementary Table S2 for details of the CI-out-closed model). Water molecules were built into the CI-out-closed, CI-out-open1 and CI-out-open2 models, which have the highest resolution for each state. Water molecules were added using the *Find Waters* command in Coot with default values and a root-mean-square deviation value starting at 6.0. Subsequent additions were performed to fill in water–water contacts, and all water molecules manually inspected for good geometry and cryoEM density. A strong density at ND4, tetrahedrally coordinated by a water molecule, an Asp and two His side chains, was modelled as a Zn^2+^ ion. The addition of non-solvent hydrogen atoms and subsequent real-space refinement, with grouped ADP refinement, and validation was performed in PHEINX v1.20.1-4887. Restraints were included for metal–protein coordination and geometry. Iteratively, models were manually adjusted in Coot to improve validation statistics after PHENIX real-space refinements.

### DDM-CI-PLs model building and refinement

The CI-in and CI-out open1 and open2 models from the as-prepared CI-PLs models were used as starting models. Water molecules were deleted, where present. For the CI-in and CI-out slack classes, a previous bovine slack model (PDB 7QSO) was used. The models were per-chain, rigid-body fit into respective maps using PHENIX. Residues, non-protein molecules and cryoEM densities were inspected and the models updated accordingly, guided by locally sharpened maps. Following Coot all-atom refinement and PHENIX real-space refinement, models were manually inspected and iterated through PHENIX real-space refinement to improve validation scores.

### Deactivated CI-PLs model building and refinement

The CI-in and CI-out open1 and open2 models from the as-prepared CI-PLs models were used as starting models and the same modelling and refinement procedures as used for the DDM-CI-PLs were implemented.

### Model validation

All model validation statistics (Supplementary Tables S1, 3–4) were generated using PHENIX, including its implementation of MolProbity^74^. Model-to-map FSC curves were generated using PHENIX.

### Structural analysis and visualisation

Map and model visualisation and image generation were performed in UCSF ChimeraX. Map regions of interest were selected with the *volume zone* command. Structural investigations on model files were also performed with UCSF ChimeraX: hydrogen bonds were detected with default parameters of the *hbonds* command; Grotthuss-competent networks were identified using the *contacts* command, with the *distanceOnly* option set to 4 Å (centre-to-centre), restricted to protonatable O and N atoms of residues D, E, H, K, S, T, and Y and water molecules only; global and per-chain comparison RMSD values were calculated with either the *matchmaker* command or the *align* command when the structures were partially superpositioned; rotation and translations of aligned models were measured with the *measure rotation* command. Cavities were detected using CASTp (1.4 Å-radius solvent probe) and visualised with CASTpyMOL 3.1 plugin and UCSF ChimeraX. The angular distributions of consensus refinements were performed in RELION, recoloured using python scripts and visualised in UCSF ChimeraX. Molleweide projections with inputs from the RELION consensus refinements were generated with a python script.

## Data availability

The structural data accession codes for the as-prepared CI-PLs dataset are: PDB 8Q48 and EMD-18141 (CI-out-closed); PDB 8Q4A and EMD-18143 (CI-out-open1); PDB 8Q49 and EMD-18142 (CI-out-open2); PDB 8Q45 and EMD-18138 (CI-in-closed); PDB 8Q47 and EMD-18140 (CI-in-open1); PDB 8Q46 and EMD-18139 (CI-in-open2). The structural data accession codes for the DDM-CI-PLs dataset are: PDB 8Q0M and EMD-18055 (CI-out-closed); PDB 8Q0O and EMD-18057 (CI-out-open2); PDB 8Q0Q and EMD-18059 (CI-out-slack); PDB 8Q0A and EMD-18051 (CI-in-closed); PDB 8Q0F and EMD-18052 (CI-in-open2); PDB 8Q0J and EMD-18054 (CI-in-slack). The structural data accession codes for the deactivated CI-PLs datasets are: PDB 8Q25 and EMD-18069 (CI-out-open1); PDB 8Q1Y and EMD-18068 (CI-out-open2); PDB 8Q1U and EMD-18067 (CI-in-open1); PDB 8Q1P and EMD-18066 (CI-in-open2). The cryoEM raw images are available from EMPIAR with the access codes EMPIAR-11678 (as-prepared CI-PLs), EMPIAR-11638 (DDM-CI-PLs), EMPIAR-11637 (deactivated CI-PLs non-tilted) and EMPIAR-11636 (deactivated CI-PLs tilted).

## Acknowledgements

We thank D. Chirgadze and S. Hardwick (University of Cambridge CryoEM facility) for assistance with grid screening and data collections. CryoEM data for the deactivated datasets were recorded at the UK National Electron Bio-Imaging Centre at the Diamond Light Source, proposal BI22238-45, funded by the Wellcome Trust, MRC and BBSRC. D.N.G. thanks the *Becoming an ‘independent’ single particle data collector* training course hosted by the Astbury Biostructure Lab (University of Leeds), eBIC (Diamond light source), Scottish Centre for Macromolecular Imaging (University of Glasgow), Midlands Regional CryoEM Facility (University of Leicester) and Birkbeck college (University of London), funded by the Wellcome Trust and Medical Research Council (218785/Z/19/Z). We thank A. J. Raine, E. E. Marcus and A. J. Nelson (MRC MBU) for IT support and I. M. Fearnley, S. Ding and N. Burger (MRC MBU) for mass spectrometry analyses. This work was supported by the Medical Research Council (MC_UU_00015/2 and MC_UU_00028/1 to J.H.).

## Author contributions

D.N.G. performed all cryoEM analyses, data processing, model building and structure analysis. J.J.W. prepared CI-PLs and performed all biochemical analyses. W.F. optimised CI purification in LMNG. Z.Y. optimised graphene oxide grid preparations with D.N.G. D.N.G. and J.H. conceived the project with input from J.J.W. J.H. supervised the project, acquired the funding and wrote the first draft of the manuscript. D.N.G. and J.J.W. produced data visualisations and wrote the final manuscript with J.H.

## Additional information

Supplementary information: The online version contains supplementary material available at [XXX]. Correspondence and requests for materials should be addressed to Judy Hirst.

[Peer review information]

Reprints and permissions information is available at www.nature.com/reprints.

## EXTENDED DATA

**Extended Data Fig. 1.**
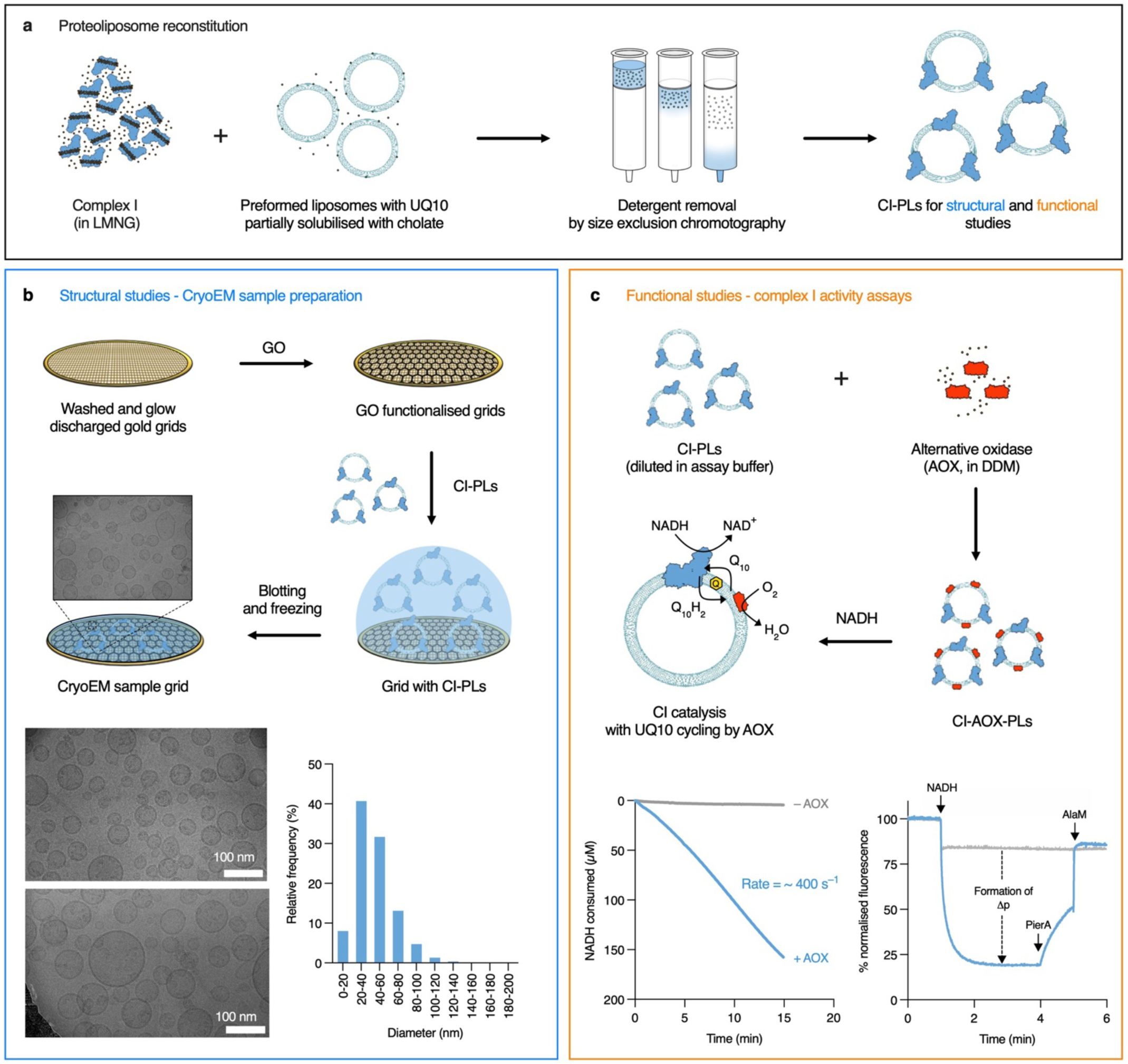
Combined biochemical, biophysical and cryoEM strategies for studying complex I-containing proteoliposomes (CI-PLs). **a)** Preparation of CI-PLs by cholate-mediated reconstitution of complex I into pre-formed liposomes. **b)** Preparation and freezing of functionalised graphene oxide (GO) grids with CI-PLs for cryoEM. Representative micrographs are shown, with the estimated size distribution determined from the edge-to-edge diameters of 1145 proteoliposomes observed in 20 randomly selected micrographs. **c)** Activation of CI-PLs for functional studies. Addition of the ubiquinol oxidase AOX (the alternative oxidase) enables studies of catalysis. In the examples shown, NADH oxidation was catalysed by 1 nM complex I, and proton pumping into the CI-PL lumen observed by quenching of the fluorescence from the probe 9-amino-6-chloro-2-methoxyacridine (ACMA), with catalysis stopped by addition of piericidin A, then Δp dissipated by addition of alamethicin.

**Extended Data Fig. 2.**
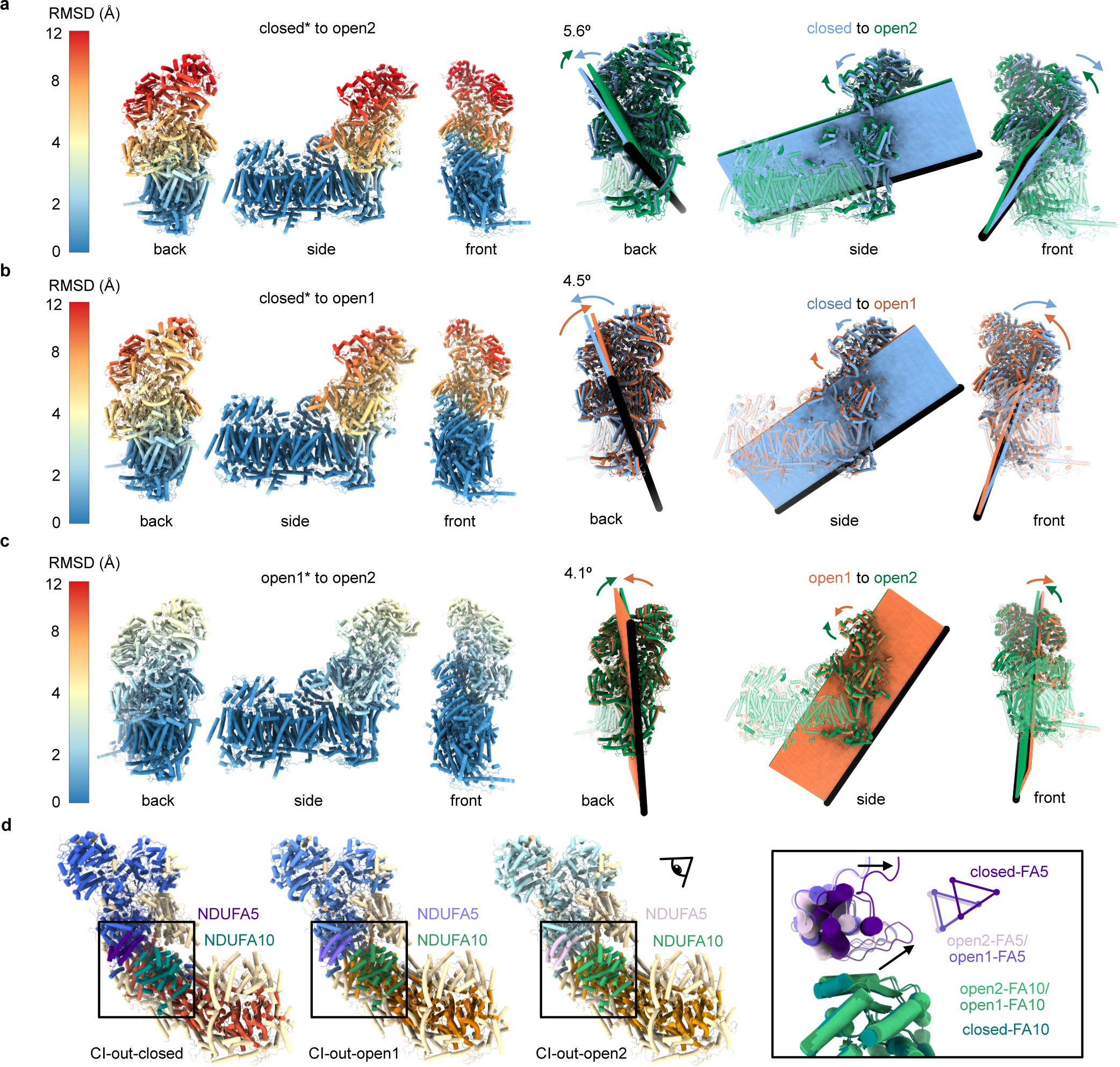
Conformational relationships between the closed, open1 and open2 states. **a)** Conformational relationships between the closed and open2 states. With the structures aligned on the membrane domain, the hydrophilic domains deviate increasingly from one another as the distance from the domain interface increases, approaching root-mean-square deviation (RMSD) values of 12 Å at the top of the domain. The RMSD values are plotted on the closed state structure (marked *). The motion can be described by rotation of the hydrophilic and heel domains (solid) about the axis of rotation (black bar) shown, while the majority of the membrane domain (translucent) does not move. The planes are shown to illustrate the angle of rotation. **b-c)** Conformational relationships between the closed and open1, and open1 and open2 states displayed as in panel **a**. The open1 state lies between the closed and open2 states, and while a similar axis of rotation can be identified between the two open states the extent of the relative displacement is much less. **d)** Variation in the relative positions of NDUFA5 (hydrophilic domain) and NDUFA10 (membrane domain) between the states. Non-NDUFA5/NDUFA10 supernumerary subunits are in wheat and core subunits in red/orange and light/dark blue for closed/open states, respectively. NDUFA5 and NDUFA10 are coloured as labelled. Boxed regions are equivalent and highlight NDUFA5/NDUFA10 interface adjustment due to rotation of the hydrophilic arm (relative to the membrane arm) between the states. For clarity, the centres of the three shifted helices are illustrated with coloured triangles also.

**Extended Data Fig. 3.**
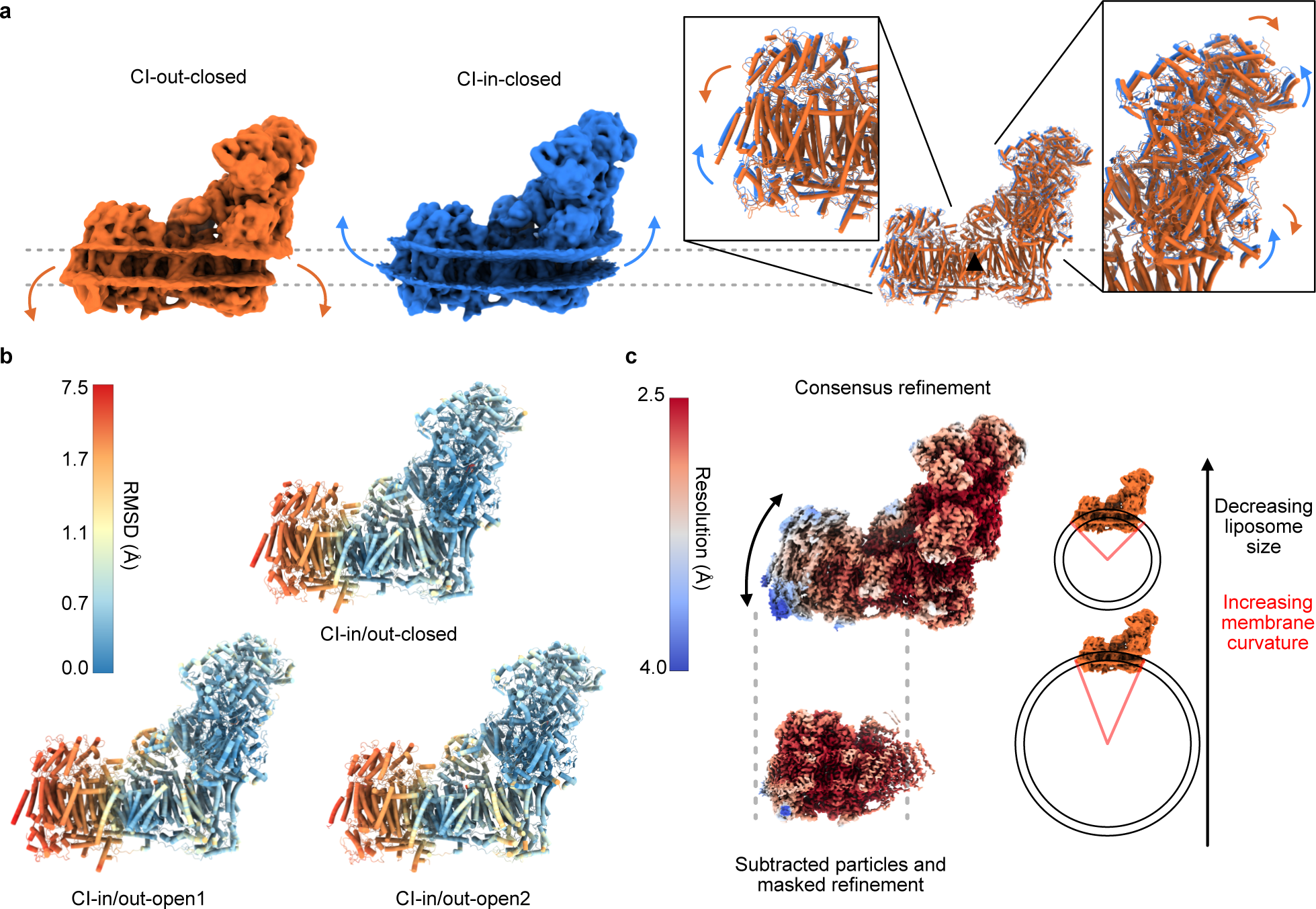
Complex I in outward-facing and inward-facing orientations and the effects of membrane curvature on its global conformation. **a)** The volumes of CI-out-closed and CI-in-closed shown with arrows to indicate the influence of membrane curvature on protein conformation, emphasised by alignment of the models on the membrane-central ND2 subunit (black triangle). Inset boxes show the extremities of the domains are affected most, with membrane-domain transmembrane helices following the curvature, and amphipathic helices at the base of the hydrophilic domain anchored at the varied membrane surface height. **b**) The effects of membrane curvature on each of the three states, shown as the root-mean-square deviation (RMSD, Å) plotted, according to the colour bar, on the respective outward models. All polypeptide chains were included in global RMSD minimization in ChimeraX before model pairwise RMSD values were calculated. **c)** Comparison of local resolutions for the distal section of the membrane domain (containing subunits ND4 and ND5) in the CI-out-closed maps from a consensus refinement with the same particles subtracted and refined on the distal section. In the consensus refinement the iron-sulphur cluster-containing hydrophilic domain dominates the global alignment, disguising the high-resolution information available for the distal section. The diagram shows how the distribution in liposome sizes creates the distribution in membrane curvature across the complex that causes the effect in the refinements.

**Extended Data Fig. 4.**
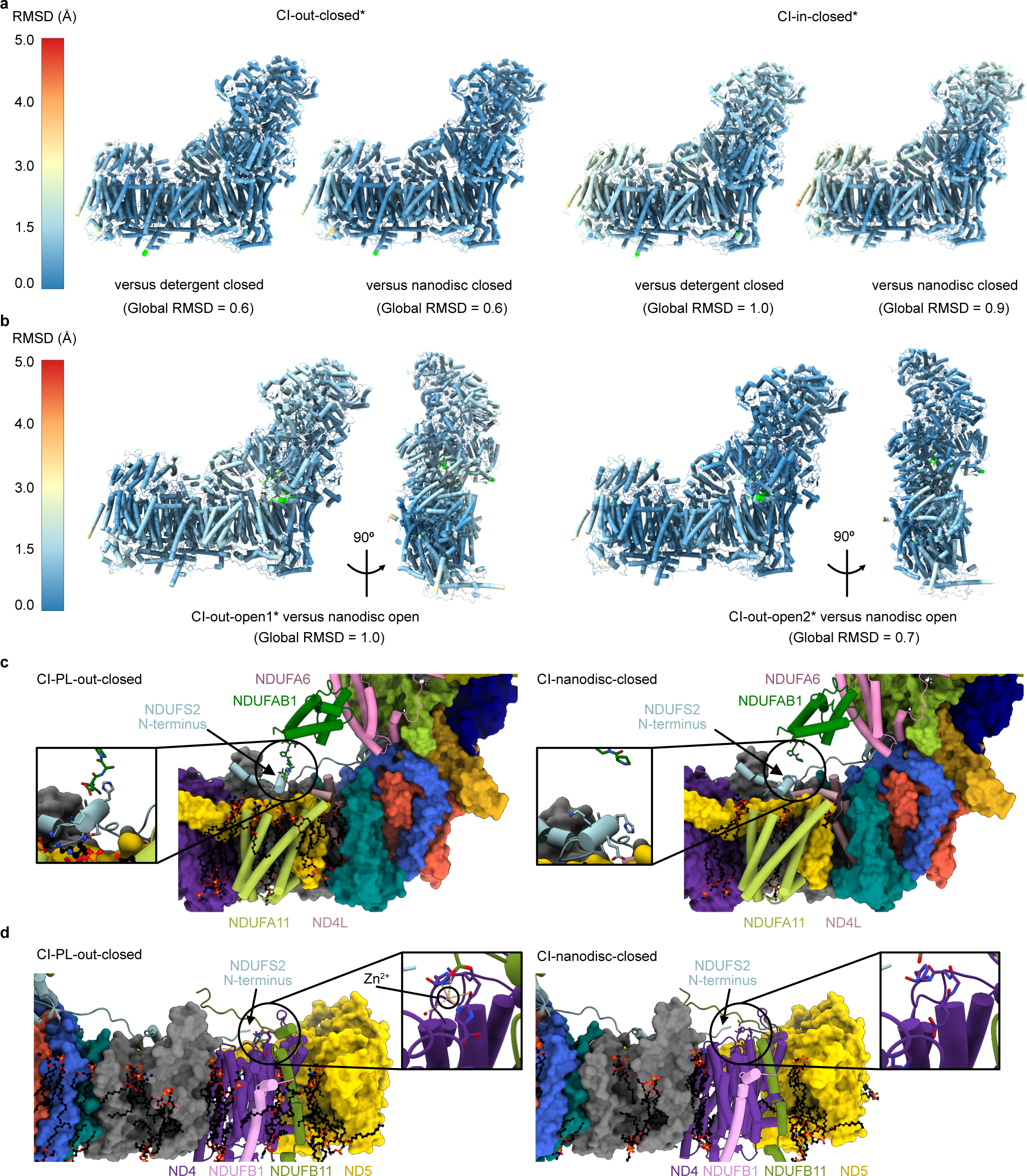
Comparisons of the closed, open1 and open2 states present in CI-PLs with equivalent structures determined in DDM micelles and nanodiscs. **a)** Root-mean-square deviation (RMSD) values (Å) between the outward/inward closed state models determined here and equivalent models determined in DDM micelles (PDB:7QSD) and nanodiscs (PDB:7QSK). The values reported in the panel titles are the global values; similarity is highest for the outward orientation, and very similar for the DDM and nanodisc models. Asterisks indicate the model used to display the values. Lime green indicates unmatched residues due to modelling differences. **b)** RMSD values (Å) between the outward open1 and open2 state models determined here and the equivalent ‘deactive’ model determined in nanodiscs (PDB:7QSN); the latter resembles open2 more than open1. **c**) The N-terminus of NDUFAB1 in complex I in CI-PLs is ordered and tethered to the membrane domain whereas in CI in nanodiscs (as well as in CI in detergent) it is disordered. **d**) The position of a Zn^2+^ modelled in the ND4 subunit in CI-PLs and the equivalent region in complex I in nanodiscs. Inset shows coordination distances of 2.3 Å for His82^ND4^ and His338^ND4^ in the axial positions of an approximately trigonal bipyramidal arrangement with Glu335^ND4^, Asp34^NDUFB1^ and a water molecule in the equatorial plane at 3.3–3.7 Å distance. For panels **c–d**, the closed state is shown in both instances, but the features shown for CI-PLs persist in the other states, with the high abundance of lipids and lateral pressure in proteoliposomes likely encouraging preservation of these features.

**Extended Data Fig. 5.**
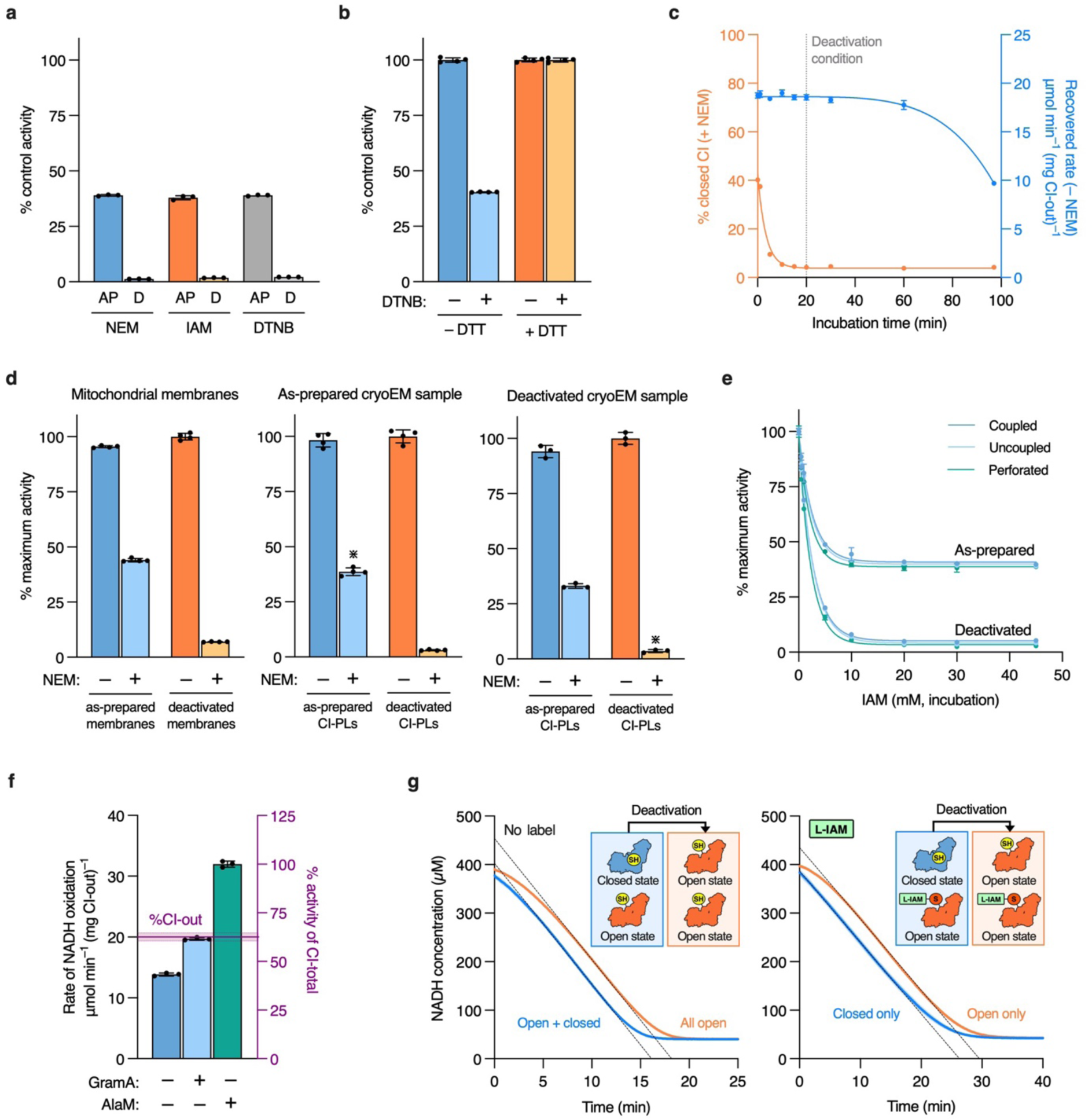
Supporting data on the biochemical characterisation and deactivation of CI-PLs. **a)** Comparison of data from three thiol-reactive reagents, N-ethyl maleimide (NEM), iodoacetamide (IAM), and 5,5′-dithiobis-(2-nitrobenzoic acid) (DTNB), used to assess the proportion of CI-closed in as-prepared (AP) and deactivated (D) CI-PLs. The proportion of the control NADH oxidation activity (AP sample incubated without the label) is reported following incubation on ice with NEM (1 mM, 30 mins), IAM (20 mM, 30 mins), or DTNB (0.2 mM, 1 hour). **b)** Treatment with DTT (1 mM, 5 mins, 20 °C) removes the DTNB from CI-open, returning the catalytic activity to the starting value (no DTNB treatment). **c)** Deactivation of CI-PLs during incubation at 37 °C. Aliquots were removed and placed on ice in the presence (orange) or absence (blue) of 1 mM NEM for 20 mins before the rate of NADH:O_2_ oxidoreduction was determined using AOX. The total recoverable rate is shown in blue and the proportion of CI-closed is shown as the NEM sensitivity in orange. The time chosen for preparation of deactive samples is indicated. **d)** NEM sensitivity measurements on the mitochondrial membranes (*n* = 4) used for cryoEM sample preparation, and on the as-prepared (*n* = 3) and deactivated CI-PLs used for cryoEM analyses. Activities are relative to the highest value in the set, and the proportion of CI-closed present under the conditions for cryoEM sample preparation (in the absence of NEM) are indicated with an asterisk. **e)** The effect of increasing IAM concentration on complex I activity in as-prepared and deactivated CI-PLs. Rates of NADH oxidation were measured with AOX under coupled, uncoupled (gramicidin A), or perforated (with alamethicin) conditions following IAM treatment. The data show IAM has derivatised both CI-out and CI-in in all cases. **f)** The rate of NADH oxidation by AOX-CI-PLs that were coupled, uncoupled (gramicidin A, CI-out only) or perforated (alamethicin, CI-out and CI-in) matches the proportion of CI-out determined by the NADH:APAD^+^ oxidoreduction assay (horizontal line, 62.6 ± 2.1%). **g)** Experiment matching that in Fig. 2d except using IAM to label the open states. The assay traces from unlabelled and IAM-labelled as-prepared CI-PLs are linear for the closed-only enzyme and display an activation lag phase when the open state is present. *n* = 3 for all experiments unless otherwise stated.

**Extended Data Fig. 6.**
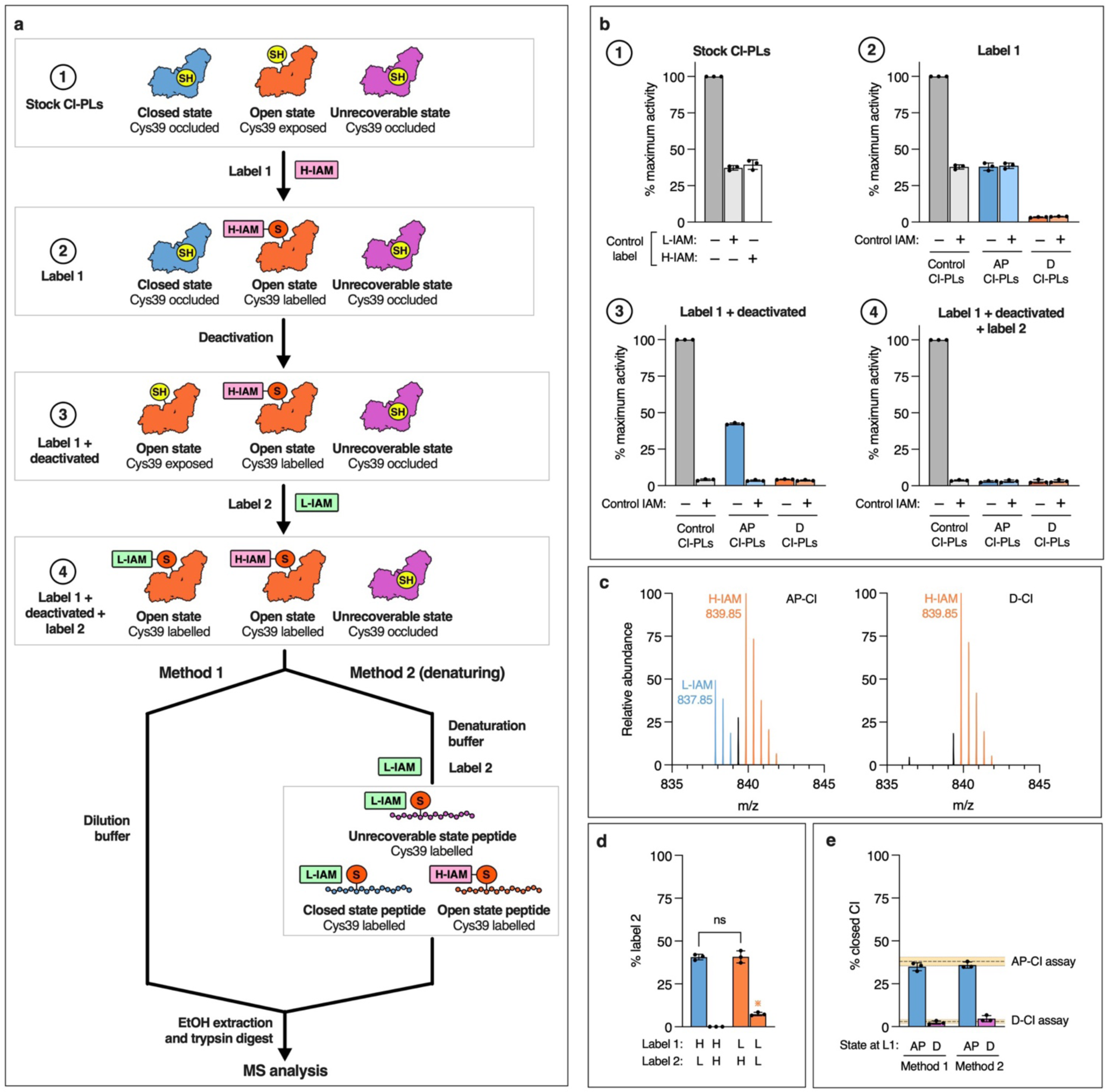
Summary of the differential isotopic labelling of CI-PLs for mass spectrometry analyses. **a)** The labelling scheme shown for a mixed population of CI-PLs containing CI-closed (Cys39^ND3^ occluded) and CI-open (Cys39^ND3^ exposed), plus a prospective population of catalytically inactive (unrecoverable) CI with Cys39 occluded (for example, aggregated enzyme). After treatment with Label 2 the sample is either treated for MS (method 1), or first denatured in the present of Label 2 (method 2) to capture the unrecoverable enzyme. **b)** NADH:O_2_ oxidoreduction assays at each step in the labelling scheme in **a** relative to the maximal activity observed with a vehicle control sample (treated identically but without label). At stage 1 the equal effect of L-IAM and H-IAM is demonstrated for an example set of as-prepared CI-PLs. For stages 2 to 4, data are presented for Control CI-PLs (treated identically but without preceding labels), as-prepared CI-PLs (AP-CI) and deactivated CI-PLs (D-CI), with the activities are shown with and without an additional IAM labelling step (± control IAM, 20 mM L-IAM, 30 mins) to confirm completion of labelling and deactivation processes. The data are mean averages (± SD) of the three independent CI-PL reconstitutions used for MS analyses. **c)** Representative mass spectra of as-prepared and deactivated CI-PLs treated according to the scheme in **a**, method 1. Peaks from the H-IAM labelled ND3 peptide (monoisotopic peak at 839.85 Da) are in orange, and those from the L-IAM labelled peptide (monoisotopic peak at 837.85 Da) are in blue. **d)** The Label-2 monoisotopic peak intensity as a proportion of the combined Label-1 and Label-2 intensities for as-prepared CI-PLs. Reversing the H-IAM and L-IAM labels does not affect the results (the percentage of CI-closed); the signal observed at 839.85 Da when L-IAM is used for both labelling steps likely arises from the natural abundance of ^13^C. The data are mean averages (± SD) of three ethanol extractions from a single sample for each condition, with statistical significance assessed by one-way ANOVA. **e)** The percentage of CI-closed determined using the scheme in **a**, method 1 and method 2 (bars) and by catalytic activity assays (horizontal lines). The data are mean averages (± SD) of three independent preparations for as-prepared and deactivated CI-PLs. *n* = 3 for all experiments.

**Extended Data Fig. 7.**
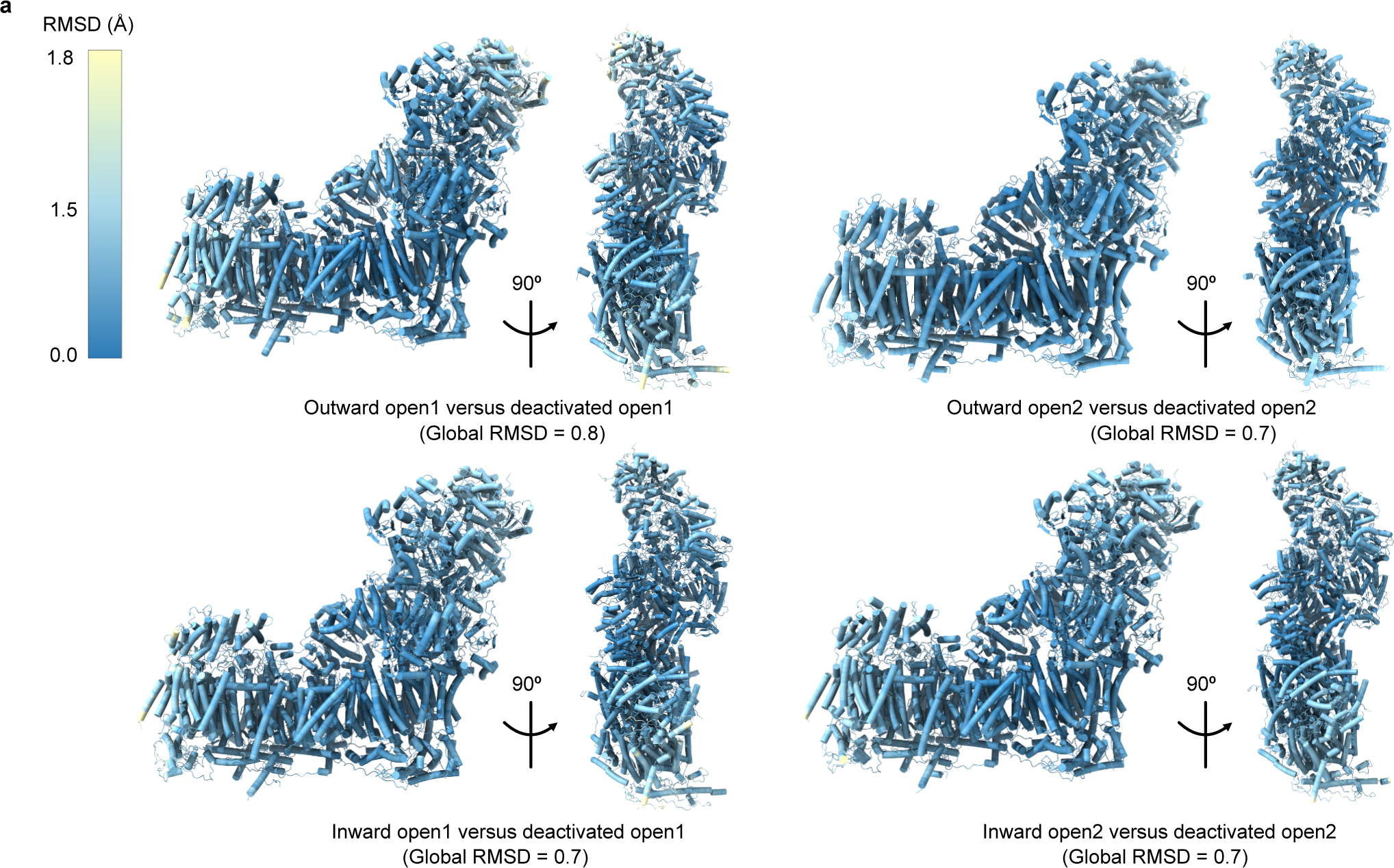
Global comparisons of the open1 and open2 states present in CI-PLs in the as-prepared and deactivated samples. Root-mean-square deviation (RMSD) values (Å) are shown for the outward- and inward-facing open1 and open2 states. The as-prepared CI-PLs models are used to display the values, according to the scale bar, and the values reported in the panel titles are the global values.

**Extended Data Fig. 8.**
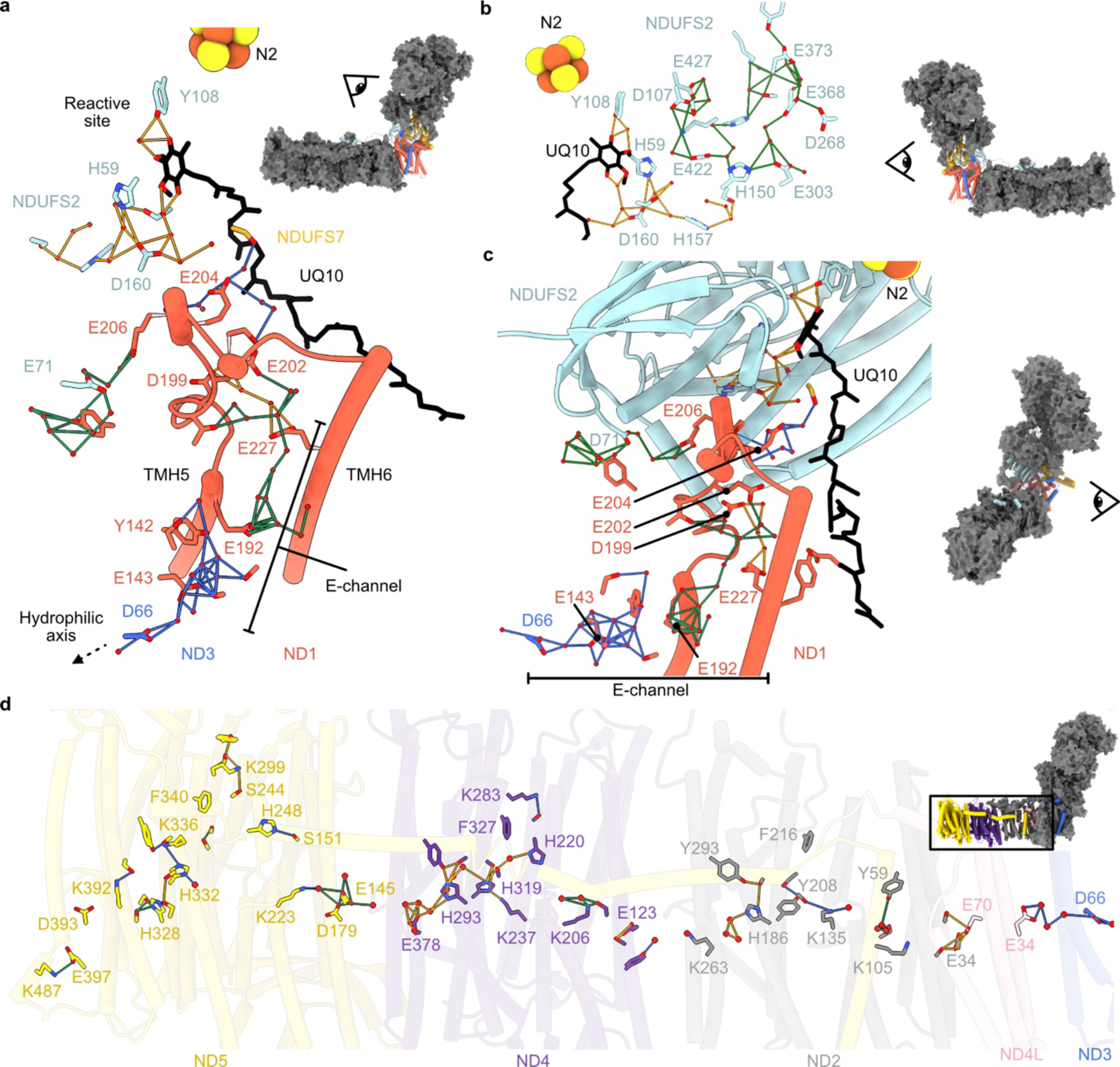
The Grotthuss networks of regions implicated in proton-transfer events in the closed state. **a**) The Grotthuss networks (4 Å sidechain oxygen/nitrogen distances) of Grotthuss-competent residues (D, E, H, K, S, Y), water molecules and the fully bound ubiquinone-10 (UQ10) are shown for the CI-out-closed state. Networks were detected in subunits ND1 (red), ND3 (dark blue), NDUFS2 (light blue) and NDUFS7 (gold), and coloured by group with minor and distant networks discarded for clarity. TMH5–6^ND1^ are shown for context. The viewpoint is indicated by the inset complex I. **b**) Networks at the reactive site of UQ10 reduction and extending through subunit NDUFS2. **c**) An alternate view of the Grotthuss networks extending from the reaction site down to the hydrophilic axis (D66^ND3^) via residues of the TMH5–6^ND1^ loop and E-channel. The acidic residues E206^ND1^, E204^ND1^, E202^ND1^, E227^ND1^, E192^ND1^ occupy positions where they may gate proton transfers between the observed networks. **d**) Grotthuss networks detected across the hydrophilic axis of the core transmembrane subunits ND3, ND4L, ND2, ND4 and ND5, with networks outside of the key residues shown removed for clarity. The apparent connectivity across the hydrophilic axis is likely due to the modest resolution in the membrane arm limiting the number of water molecules modelled.

**Extended Data Fig. 9.**
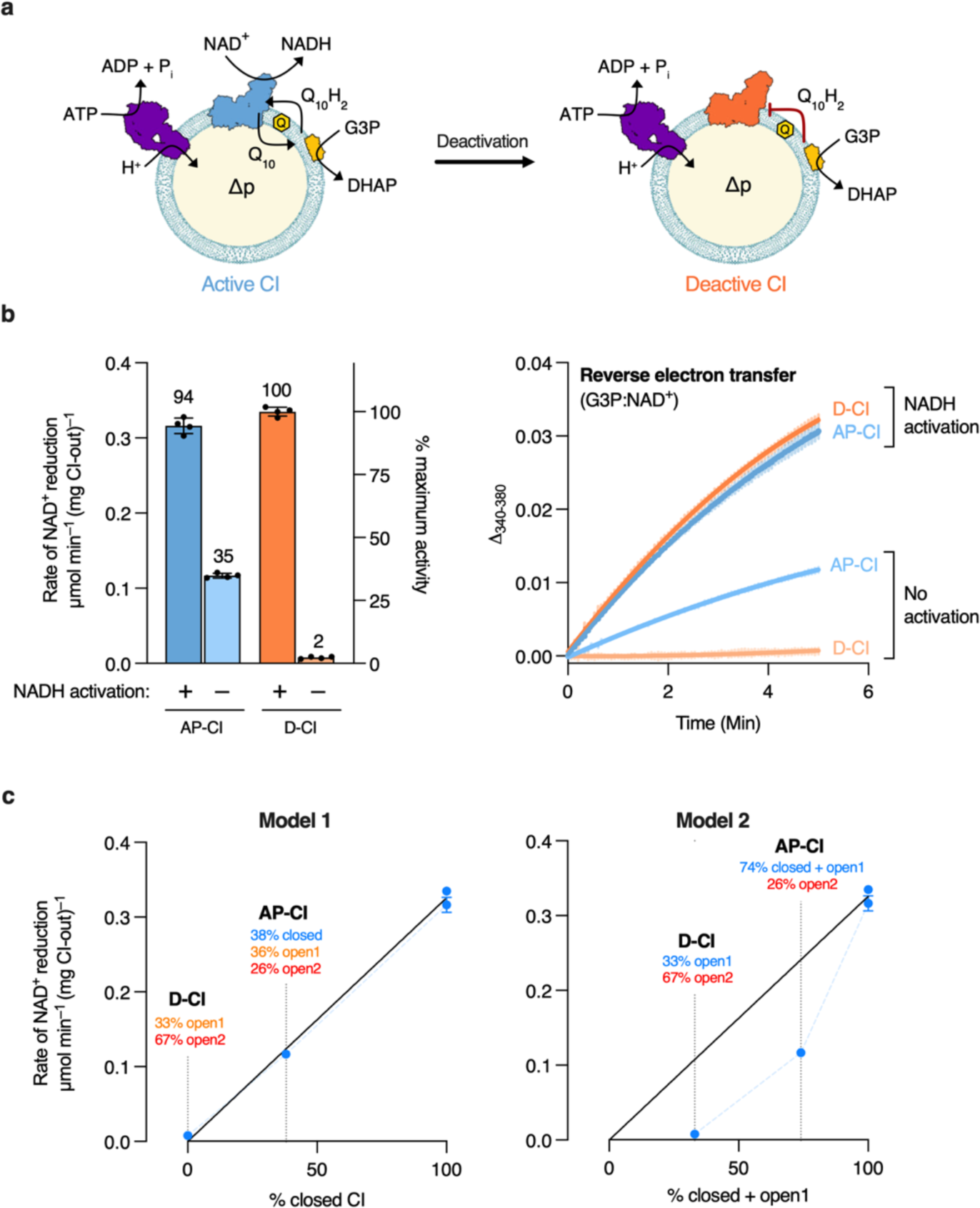
Assessment of the biochemical state of complex I in proteoliposomes by reverse electron transfer. **a)** Schematic representation of co-reconstituted ATPase-CI-PLs for reverse electron transfer. The PLs are supplemented with glycerol 3-phosphate dehydrogenase (GlpD) to reduce the UQ10 pool (the oxidised and reduced states are denoted Q_10_ and Q_10_H_2_, respectively), and the proton-motive force (Δp) is generated by ATP hydrolysis. In the active state CI catalyses RET (observed as reduction of NAD^+^ to NADH), while in the deactive state it is incapable of RET catalysis. **b)** NAD^+^ reduction by as-prepared (AP) and deactivated (D) ATPase-CI-PLs. Samples pre-activated by addition of NADH are indicated (see methods for the activation procedure). The rates of catalysis, relative to the maximum observed activity, are given alongside the assay traces for NAD^+^ reduction. **c)** Correlation of the proportions of structural states observed by cryoEM to experimentally determined rates of NAD^+^ reduction. Model 1 assumes that only CI-closed is able to catalyse RET; Model 2 assumes that both CI-closed and CI-open1 are able to catalyse RET.

**Extended Data Table 1.** The status of structural features that differ between the closed and open states in mammalian complex I presented for models of complex I from *Bos taurus* determined here. The ‘active’ (A) and ‘deactive’ (D) indicators used are related to those described for the resting active (PDB 7QSK) and deactive (PDB 7QSN) states of bovine complex I in nanodiscs. The nanodisc slack state (7QSO) is included for reference. **ND6-TMH3:** A, α-helical or D, π-bulge at F68/Y70. **ND3 TMH1–2**, **ND1 TMH5–6 and NDUFS2 β1–β2 loops:** A, ordered in active nanodisc conformation; A-like, A with only minor differences; Alt, ordered but in a noticeably alternate backbone conformation; D-like, β-strands in NDUFS2 ordered but loop disordered; D, disordered. **ND1 TMH4:** A, bent helix with Y127-Cα in the A position; D, straight helix with Y127-Cα in the deactive nanodisc position. **NDUFS7 Arg77 loop:** A, Arg pointing away from NDUFS2 and an adjacent loop; Mixed, Arg pointing away from NDUFS2 but an adjacent β-strand; D, Arg pointing towards NDUFS2 and an adjacent β-strand.

| Preparation and state name | PDB/EMDB | Resolution | NDUFA5/10 interface (Å<sup>2</sup>) [residues buried] | ND6 TMH3 | ND3 TMH1–2 loop | ND1 TMH 5–6 loop | ND1 TMH4 | NDUFS2 β1–β2 loop | NDUFS7 Arg77 loop |
| --- | --- | --- | --- | --- | --- | --- | --- | --- | --- |
| CI-out-closed | 8Q48/18141 | 2.5 | 361 [20] | A | A | A | A | A | A |
| CI-in-closed | 8Q45/18138 | 2.7 | 393 [20] | A | A | A | A | A | A |
| CI-out-open1 | 8Q4A/18143 | 2.6 | 127 [14] | D | Alt | A-like | D | D-like | D |
| CI-in-open1 | 8Q47/18140 | 2.9 | 163 [15] | D | Alt | A-like | D | D-like | D |
| CI-out-open2 | 8Q49/18142 | 2.6 | 126 [9] | D | D | A-like | D | D-like | D |
| CI-in-open2 | 8Q46/18139 | 2.6 | 158 [15] | D | D | A-like | D | D-like | D |
| DDM CI-out-closed | 8Q0M/18055 | 3.1 | 301 [18] | A | A | A | A | A | A |
| DDM CI-in-closed | 8Q0A/18051 | 3.1 | 338 [17] | A | A | A | A | A | A |
| DDM CI-out-open2 | 8Q0O/18057 | 3.1 | 119 [8] | D | D | A-like | D | D-like | D |
| DDM CI-in-open2 | 8Q0F/18052 | 3.1 | 156 [14] | D | D | A-like | D | D-like | D |
| DDM CI-out-slack | 8Q0Q/18059 | 3.6 | 76 [8] | D | D | Alt | D | Alt | Mixed |
| DDM CI-in-slack | 8Q0J/18054 | 3.8 | 96 [7] | D | D | Alt | D | Alt | Mixed |
| Deactivated CI-out-open1 | 8Q25/18069 | 2.8 | 80 [7] | D | Alt | A-like | D | D-like | D |
| Deactivated CI-in-open1 | 8Q1U/18067 | 3.3 | 168 [15] | D | Alt | A-like | D | D-like | D |
| Deactivated CI-out-open2 | 8Q1Y/18068 | 2.6 | 124 [9] | D | D | A-like | D | D-like | D |
| Deactivated CI-in-open2 | 8Q1P/18066 | 2.9 | 154 [15] | D | D | A-like | D | D-like | D |
| Nanodisc CI-closed | 7QSK/14132 | 2.8 | 354 [19] | A | A | A | A | A | A |
| Nanodisc CI-open | 7QSN/14139 | 2.8 | 129 [9] | D | D | D | D | D | D |
| Nanodisc CI-slack | 7QSO/14140 | 3.0 | 99 [10] | D | D | Alt | D | Alt | Mixed |

