## Supplementary Information for "Structural basis of a regulatory switch in mammalian complex I"

### SUPPLEMENTARY DATA

#### Table S1 | CryoEM data collection, refinement and validation statistics for the CI-PLs dataset.

| | CI-PLs dataset (purified in LMNG, as-prepared) | | | | | |
|---|---|---|---|---|---|---|
| **Data collection and processing** | | | | | | |
| Nominal magnification | 81,000 | | | | | |
| Voltage (kV) | 300 | | | | | |
| Electron exposure ($e^{–}$ Å$^{-2}$) | 45 | | | | | |
| Targeted defocus range (µm) | -0.9 to -2.3 | | | | | |
| Calibrated pixel size (Å) | 1.066 | | | | | |
| Symmetry imposed | *C*1 | | | | | |
| EMPIAR code | EMPIAR-11678 | | | | | |
| Initial particle images (no.) | 5,688,318 | | | | | |
| State and model (PDB/EMDB codes) | **CI-out-closed (8Q48/18141)** | **CI-out-open1 (8Q4A/18143)** | **CI-out-open2 (8Q49/18142)** | **CI-in-closed (8Q45/18138)** | **CI-in-open1 (8Q47/18140)** | **CI-in-open2 (8Q46/18139)** |
| Final particle images (no.) | 183,021 | 175,986 | 127,085 | 63,569 | 30,701 | 133,884 |
| Map resolution (Å) (FSC 0.143) | 2.5 | 2.6 | 2.6 | 2.7 | 2.9 | 2.6 |
| Map resolution range (Å) | 2.3–5.5 | 2.3–5.6 | 2.4–6.6 | 2.5–5.8 | 2.6–6.6 | 2.3–5.3 |
| Map sharpening (RELION) *B* factor (Å$^2$) | −27 | −33 | −33 | −28 | −28 | −31 |
| **Refinement** | | | | | | |
| Initial model used | 7QSK | 7QSN | 7QSN | 7QSK | 7QSN | 7QSN |
| Model resolution (Å) (FSC 0.5) | 2.6 | 2.7 | 2.8 | 2.7 | 2.9 | 2.7 |
| Model composition | | | | | | |
| Nonhydrogen atoms | 71,834 | 70,996 | 70,056 | 71,363 | 70,199 | 68,995 |
| Protein residues | 8286 | 8272 | 8243 | 8293 | 8274 | 8243 |
| Ligands | 91 | 83 | 81 | 86 | 74 | 71 |
| Waters | 1122 | 953 | 732 | n/a | n/a | n/a |
| *B* factors mean (Å$^2$) | | | | | | |
| Protein | 49.10 | 52.81 | 47.08 | 46.96 | 50.13 | 48.53 |
| Ligand | 48.65 | 52.44 | 50.51 | 48.77 | 51.10 | 49.69 |
| Water | 35.38 | 39.19 | 39.33 | n/a | n/a | n/a |
| RMS deviations | | | | | | |
| Bond lengths (Å) | 0.003 | 0.004 | 0.003 | 0.002 | 0.004 | 0.004 |
| Bond angles (°) | 0.511 | 0.503 | 0.474 | 0.502 | 0.493 | 0.514 |
| Validation | | | | | | |
| MolProbity score | 1.29 | 1.57 | 1.68 | 1.53 | 1.77 | 1.58 |
| Clashscore | 3.72 | 3.68 | 3.92 | 3.17 | 4.32 | 3.41 |
| Rotamer outliers (%) | 1.12 | 1.85 | 2.30 | 2.63 | 2.75 | 2.56 |
| Cß outliers (%) | 0.00 | 0.00 | 0.00 | 0.00 | 0.00 | 0.00 |
| Ramachandran plot | | | | | | |
| Favored (%) | 97.57 | 96.73 | 96.50 | 97.47 | 96.54 | 97.24 |
| Allowed (%) | 2.42 | 3.26 | 3.47 | 2.53 | 3.42 | 2.72 |
| Outliers (%) | 0.01 | 0.01 | 0.02 | 0.00 | 0.04 | 0.04 |
| Rama-Z (Ramachandran plot Z-score, RMSD) | | | | | | |
| Whole | 0.18 (0.09) | 0.08 (0.10) | 0.06 (0.10) | 0.05 (0.10) | -0.35 (0.10) | -0.23 (0.09) |
| Helix | 0.36 (0.08) | 0.38 (0.08) | 0.50 (0.08) | 0.30 (0.08) | 0.10 (0.08) | 0.06 (0.08) |
| Sheet | 0.03 (0.25) | -0.06 (0.25) | -0.29 (0.27) | -0.14 (0.26) | -0.33 (0.26) | -0.36 (0.26) |
| Loop | 0.02 (0.11) | -0.18 (0.11) | -0.34 (0.11) | -0.10 (0.11) | -0.46 (0.11) | -0.20 (0.11) |

**Table S2 | Summary of modifications and cofactors of the model for CI-out-closed (PDB: 8Q48).**

| Subunit | Alternative names | Chain | Total residues | N-terminal modification | Modelled residues (%) | Modelled cofactors and modifications |
|---|---|---|---|---|---|---|
| NDUFV1 | 51 kDa, Nqo1, NuoF | F | 444 | Δ 1–20 | 7–438 (97.3) | FMN, 4Fe4S |
| NDUFV2 | 24 kDa, Nqo2, NuoE | E | 217 | Δ 1–32 | 4–217 (98.6) | 2Fe2S |
| NDUFS1 | 75 kDa, Nqo3, NuoG | G | 704 | Δ 1–23 | 6–693 (97.7) | 2Fe2S, 2 x 4Fe4S |
| NDUFS2 | 49 kDa, Nqo4, NuoC | D | 430 | Δ 1–33 | 1–430 (100) | Dimethyl-Arg85 |
| NDUFS3 | 30 kDa, Nqo5, NuoC | C | 228 | Δ 1–38 | 6–214 (91.6) | |
| NDUFS7 | PSST, Nqo6, NuoB | B | 179 | Δ 1–37 | 25–179 (86.6) | 4Fe4S |
| NDUFS8 | TYKY, Nqo9, NuoI | I | 176 | Δ 1–36 | 1–176 (100) | 2 x 4Fe4S |
| ND1 | Nqo8, NuoH | H | 318 | N-formyl | 1–318 (100) | N-formyl |
| ND2 | Nqo14, NuoN | N | 347 | N-formyl | 1–347 (100) | N-formyl |
| ND3 | Nqo7, NuoA | A | 115 | N-formyl | 1–115 (100) | N-formyl |
| ND4 | Nqo13, NuoM | M | 459 | N-formyl | 1–459 (100) | N-formyl |
| ND4L | Nqo11, NuoK | K | 98 | N-formyl | 1–98 (100) | N-formyl |
| ND5 | Nqo12, NuoL | L | 606 | N-formyl | 1–606 (100) | N-formyl |
| ND6 | Nqo10, NuoJ | J | 175 | N-formyl | 1–175 (100) | N-formyl |
| NDUFV3 | 10 kDa | s | 75 | Δ 1–34 | 31–75 (60.0) | |
| NDUFS4 | 18 kDa | Q | 133 | Δ 1–42 | 5–133 (97.0) | |
| NDUFS5 | 15 kDa | e | 105 | –Met | 1–99 (94.3) | 2 x Cys–Cys |
| NDUFS6 | 13 kDa | R | 96 | Δ 1–28 | 1–96 (100) | $Zn^{2+}$ |
| NDUFA1 | MWFE | a | 70 | – | 1–70 (100) | |
| NDUFA2 | B8 | S | 98 | –Met +Ac | 12–98 (88.8) | |
| NDUFA3 | B9 | b | 83 | –Met +Ac | 1–83 (100) | N-acetyl |
| NDUFA5 | B13 | V | 115 | –Met +Ac | 1–115 (99.1) | |
| NDUFA6 | B14 | W | 127 | –Met +Ac | 13–127 (90.6) | |
| NDUFA7 | B14.5a | r | 112 | –Met +Ac | 1–72, 90–112 (84.8) | N-acetyl |
| NDUFA8 | PGIV | X | 171 | –Met | 1–171 (100) | |
| NDUFA9 | 39 kDa | P | 345 | Δ 1–35 | 1–342 (99.1) | NADPH |
| NDUFA10 | 42 kDa | O | 320 | Δ 1–23 | 1–320 (100) | $Mg^{2+}$–dGTP |
| NDUFA11 | B14.7 | Y | 140 | –Met +Ac | 1–140 (100) | 2 x Cys–Cys, N-acetyl |
| NDUFA12 | B17.2 | q | 145 | +Ac | 1–145 (100) | N-acetyl |
| NDUFA13 | B16.6 | Z | 143 | –Met +Ac | 3–143 (98.6) | |
| NDUFAB1α | SDAPα | T | 88 | Δ 1–46 | 1–88 (100) | 4'-phosphopantethine + 3-hydroxytetradecanoate |
| NDUFAB1β | SDAPβ | U | | | | |
| NDUFB1 | MNLL | f | 57 | –Met | 1–57 (100) | |
| NDUFB2 | AGGG | j | 72 | Δ 1–36 | 1–71 (98.6) | |
| NDUFB3 | B12 | k | 97 | –Met +Ac partial | 1–81 (83.5) | |
| NDUFB4 | B15 | m | 128 | –Met +Ac | 1–128 (100) | N-acetyl |
| NDUFB5 | SGDH | h | 143 | Δ 1–46 | 6–143 (96.5) | |
| NDUFB6 | B17 | i | 127 | –Met +Ac | 1–127 (100) | N-acetyl |
| NDUFB7 | B18 | o | 136 | –Met +Myr | 1–122 (89.7) | N-myristoyl |
| NDUFB8 | ASHI | l | 158 | Δ 1–28 | 3–158 (98.7) | |
| NDUFB9 | B22 | n | 178 | –Met +Ac | 8–178 (96.1) | |
| NDUFB10 | PDSW | p | 175 | –Met | 2–175 (99.4) | |
| NDUFB11 | ESSS | g | 125 | Δ 1–29 | 22–122 (80.8) | |
| NDUFC1 | KFYI | c | 49 | Δ 1–27 | 1–49 (100) | |
| NDUFC2 | B14.5b | d | 120 | +Ac partial | 1–120 (100) | N-acetyl |

**Table S3 | CryoEM data collection, refinement and validation statistics for the DDM-CI-PLs dataset.**

| | DDM-CI-PLs dataset (purified in DDM, as-prepared) | | | | | |
|---|---|---|---|---|---|---|
| **Data collection and processing** | | | | | | |
| Nominal magnification | 81,000 | | | | | |
| Voltage (kV) | 300 | | | | | |
| Electron exposure ($e^{-}$ Å$^{-2}$) | 40 | | | | | |
| Targeted defocus range (μm) | -1.0 to -3.0 | | | | | |
| Calibrated pixel size (Å) | 1.35 | | | | | |
| Symmetry imposed | *C*1 | | | | | |
| EMPIAR code | EMPIAR-11638 | | | | | |
| Initial particle images (no.) | 2,313,863 | | | | | |
| State and model (PDB/EMDB codes) | **CI-out-closed (8Q0M/18055)** | **CI-out-open2 (8Q0O/18057)** | **CI-out-slack (8Q0Q/18059)** | **CI-in-closed (8Q0A/18051)** | **CI-in-open2 (8Q0F/18052)** | **CI-in-slack (8Q0J/18054)** |
| Final particle images (no.) | 81,459 | 123,011 | 45,544 | 68,095 | 116,474 | 45,661 |
| Map resolution (Å) (FSC 0.143) | 3.1 | 3.1 | 3.6 | 3.1 | 3.1 | 3.8 |
| Map resolution range (Å) | 2.9–6.6 | 2.9–6.4 | 3.2–9.3 | 2.8–6.6 | 2.9–6.6 | 3.4–9.8 |
| Map sharpening (RELION) *B* factor (Å$^2$) | −41 | −49 | −53 | −35 | −49 | −52 |
| **Refinement** | | | | | | |
| Initial model used | 8Q48 | 8Q49 | 7QSO | 8Q45 | 8Q46 | 7QSO |
| Model resolution (Å) (FSC 0.5) | 3.1 | 3.2 | 3.6 | 3.1 | 3.2 | 3.8 |
| Model composition | | | | | | |
| Nonhydrogen atoms | 71,676 | 70,323 | 65,867 | 70,959 | 70,023 | 65,999 |
| Protein residues | 8286 | 8243 | 8004 | 8286 | 8243 | 8004 |
| Ligands | 88 | 77 | 37 | 80 | 68 | 37 |
| Waters | n/a | n/a | n/a | n/a | n/a | n/a |
| *B* factors mean (Å$^2$) | | | | | | |
| Protein | 54.80 | 48.26 | 83.22 | 52.04 | 56.43 | 93.64 |
| Ligand | 55.45 | 46.54 | 80.27 | 52.81 | 55.50 | 87.57 |
| Water | n/a | n/a | n/a | n/a | n/a | n/a |
| RMS deviations | | | | | | |
| Bond lengths (Å) | 0.003 | 0.004 | 0.005 | 0.005 | 0.003 | 0.004 |
| Bond angles (°) | 0.491 | 0.505 | 0.528 | 0.517 | 0.486 | 0.496 |
| Validation | | | | | | |
| MolProbity score | 1.41 | 1.83 | 2.15 | 1.70 | 1.76 | 2.20 |
| Clashscore | 3.46 | 4.43 | 8.21 | 3.64 | 4.18 | 8.58 |
| Rotamer outliers (%) | 1.50 | 2.60 | 2.28 | 2.41 | 2.69 | 2.49 |
| Cß outliers (%) | 0.00 | 0.00 | 0.00 | 0.00 | 0.00 | 0.00 |
| Ramachandran plot | | | | | | |
| Favored (%) | 97.27 | 95.72 | 93.37 | 96.23 | 96.42 | 93.30 |
| Allowed (%) | 2.71 | 4.25 | 6.62 | 3.75 | 3.53 | 6.64 |
| Outliers (%) | 0.01 | 0.04 | 0.01 | 0.01 | 0.05 | 0.05 |
| Rama-Z (Ramachandran plot Z-score, RMSD) | | | | | | |
| Whole | 0.44 (0.10) | -0.43 (0.09) | -0.93 (0.09) | -0.11 (0.09) | -0.03 (0.10) | -0.75 (0.10) |
| Helix | 0.69 (0.08) | 0.12 (0.08) | -0.23 (0.08) | 0.26 (0.08) | 0.38 (0.08) | 0.01 (0.09) |
| Sheet | -0.10 (0.26) | -0.67 (0.27) | -0.92 (0.29) | -0.53 (0.25) | -0.70 (0.27) | -1.29 (0.27) |
| Loop | 0.00 (0.11) | -0.61 (0.11) | -0.89 (0.10) | -0.26 (0.11) | -0.28 (0.11) | -0.84 (0.10) |

**Table S4 | CryoEM data collection for the deactivated tilted and non-tilted CI-PLs datasets, and the refinement and validation statistics for the four states from the combined deactivated CI-PLs dataset.**

| | Deactivated CI-PLs dataset (non-tilted) | Deactivated CI-PLs dataset (tilted 20º) |
|---|---|---|
| **Data collection and processing** | | |
| Nominal magnification | 81,000 | 81,000 |
| Voltage (kV) | 300 | 300 |
| Electron exposure (e⁻ Å$^{-2}$) | 40 | 40 |
| Targeted defocus range (μm) | -0.9 to -2.3 | -0.8 to -2.0 |
| Calibrated pixel size (Å) | 1.072 | 1.072 |
| Symmetry imposed | *C*1 | *C*1 |
| EMPIAR code | EMPIAR-11637 | EMPIAR-11636 |
| Initial particle images (no.) | 932,186 | 878,425 |

| | CI-out-open1 | CI-out-open2 | CI-in-open1 | CI-in-open2 |
|---|---|---|---|---|
| PDB/EMDB codes (combined datasets) | **8Q25/18069** | **8Q1Y/18068** | **8Q1U/18067** | **8Q1P/18066** |
| Final particle images (no.) | 45,912 | 93,255 | 14,905 | 58,386 |
| Map resolution (Å) (FSC 0.143) | 2.8 | 2.6 | 3.3 | 2.9 |
| Map resolution range (Å) | 2.4–7.2 | 2.3–6.7 | 2.9–8.2 | 2.5–6.5 |
| Map sharpening (RELION) *B* factor (Å$^2$) | −26 | −25 | −36 | −33 |
| **Refinement** | | | | |
| Initial model used | 8Q4A | 8Q49 | 8Q47 | 8Q46 |
| Model resolution (Å) (FSC 0.5) | 2.8 | 2.7 | 3.2 | 2.9 |
| Model composition | | | | |
| Nonhydrogen atoms | 70,996 | 70,571 | 70,058 | 70,162 |
| Protein residues | 8272 | 8243 | 8274 | 8243 |
| Ligands | 83 | 81 | 72 | 71 |
| Waters | n/a | n/a | n/a | n/a |
| *B* factors mean (Å$^2$) | | | | |
| Protein | 56.54 | 45.47 | 48.11 | 50.13 |
| Ligand | 55.07 | 50.32 | 51.69 | 53.24 |
| Water | n/a | n/a | n/a | n/a |
| RMS deviations | | | | |
| Bond lengths (Å) | 0.003 | 0.004 | 0.004 | 0.003 |
| Bond angles (°) | 0.507 | 0.477 | 0.500 | 0.475 |
| Validation | | | | |
| MolProbity score | 1.54 | 1.58 | 1.84 | 1.54 |
| Clashscore | 3.40 | 3.11 | 4.31 | 3.06 |
| Rotamer outliers (%) | 2.01 | 2.52 | 2.61 | 2.55 |
| Cß outliers (%) | 0.00 | 0.00 | 0.00 | 0.00 |
| Ramachandran plot | | | | |
| Favored (%) | 97.00 | 97.00 | 95.51 | 97.32 |
| Allowed (%) | 2.99 | 2.99 | 4.45 | 2.66 |
| Outliers (%) | 0.01 | 0.01 | 0.04 | 0.02 |
| Rama-Z (Ramachandran plot Z-score, RMSD) | | | | |
| Whole | -0.48 (0.09) | -0.16 (0.09) | -0.58 (0.09) | -0.10 (0.09) |
| Helix | -0.09 (0.08) | 0.26 (0.08) | 0.04 (0.08) | 0.20 (0.08) |
| Sheet | -0.67 (0.25) | -0.18 (0.27) | -0.69 (0.27) | -0.08 (0.27) |
| Loop | -0.36 (0.11) | -0.40 (0.11) | -0.73 (0.10) | -0.23 (0.11) |

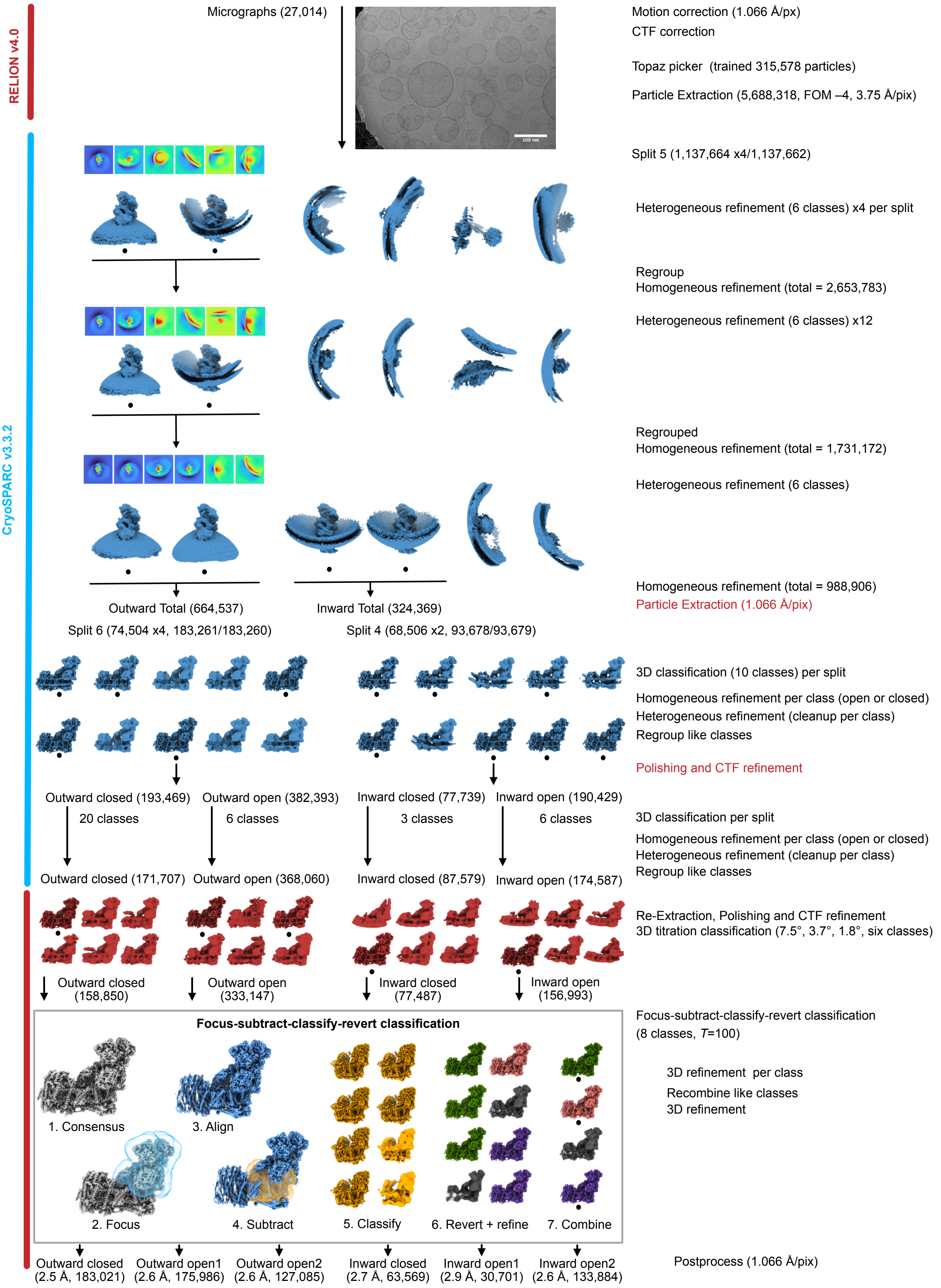


**Fig. S1 | Classification scheme for the CI-PLs dataset (purified in LMNG, as prepared).** An example micrograph and 3D volumes are shown along with cryoEM density heatmaps for volumes in cryoSPARC heterogeneous refinements. Volumes taken forward are indicated with a black dot. The software used to perform each step is indicated by the left-hand colour bar (red text indicates RELION-specific stage exceptions). The focus-subtract-classify-revert classification method is outlined in the grey box using example volumes for the outward closed analysis.

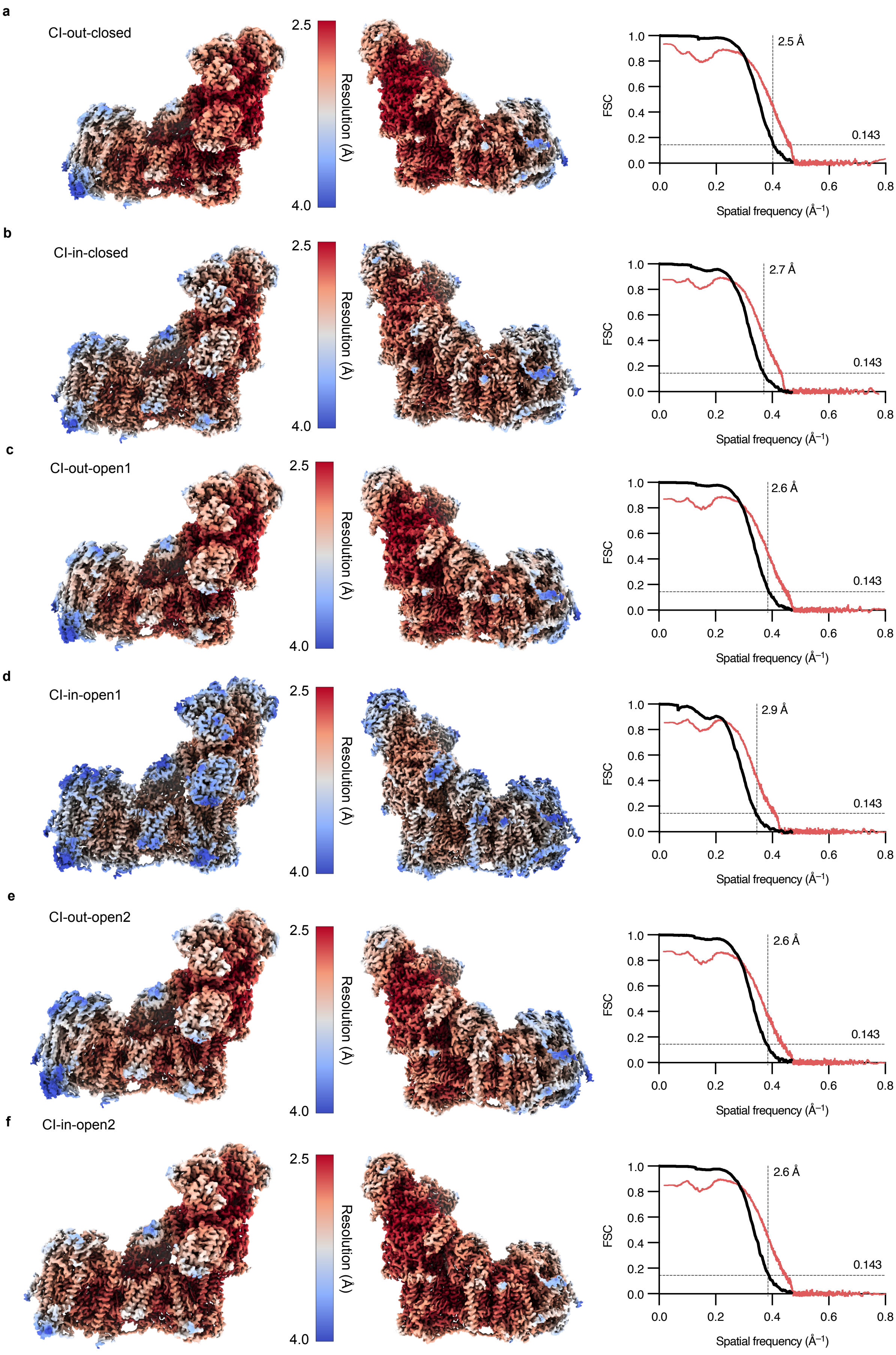


**Fig. S2 | Global and local resolution of the consensus classes of the CI-PLs dataset (purified in LMNG, as prepared).** The final classes **a**) CI-out-closed, **b**) CI-in-closed, **c**) CI-out-open1, **d**) CI-in-open1, **e**) CI-out-open2, and **f**) CI-in-open2 were analysed for local resolution distribution using RELION v4.0-beta. Values are plotted according to the colour key. Global Fourier Shell Correlation (FSC) scores (cutoff = 0.143) calculated in RELION v4.0-beta are plotted (black) along with model–map FSCs (red), calculated using the validation suite in PHENIX v1.20.1-4487.

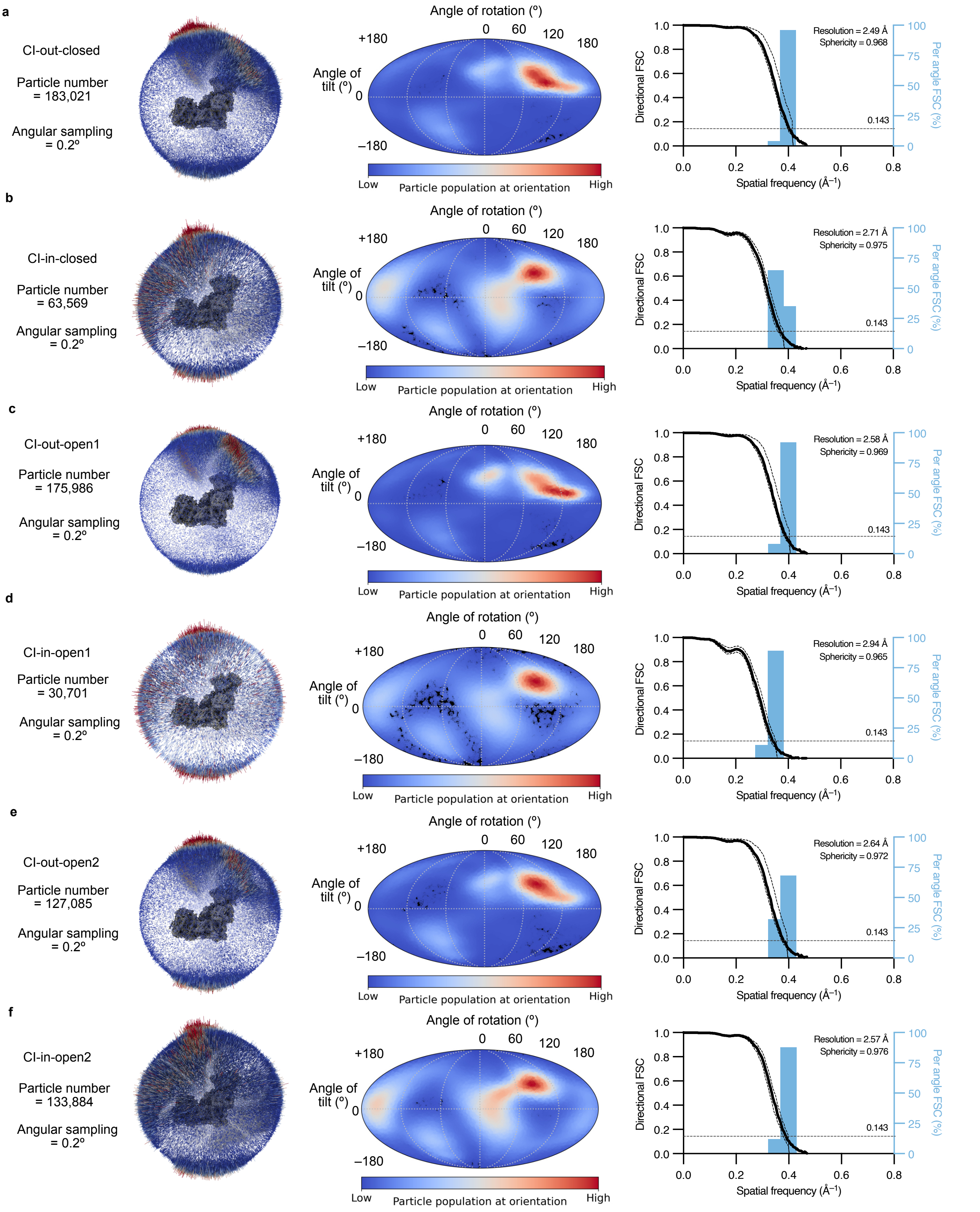


**Fig. S3 | Orientation analyses of the consensus classes of the CI-PLs dataset (purified in LMNG, as prepared).** The final classes **a**) CI-out-closed, **b**) CI-in-closed, **c**) CI-out-open1, **d**) CI-in-open1, **e**) CI-out-open2, and **f**) CI-in-open2 were inspected for the orientation distribution of the particles in the final consensus alignment using 3D angular and Mollweide plots. Equivalent data are shown in each case. The orientation distribution in the Mollweide plots is relative to the input reference volume, so the complex I 3D volumes (grey) in each case are shown in the 3D angular distribution spheres for reference. The assessment of uniformity of resolution in each direction (sphericity, where 1 is completely uniform) was performed by dividing the sphere up into cones and calculating directional Fourier-shell correlations (FSCs), plotted as histograms, using the 3DFSC Program Suite v3.0. The 3DFSC global FSC curve is shown in black (± S.D. of directional FSCs).

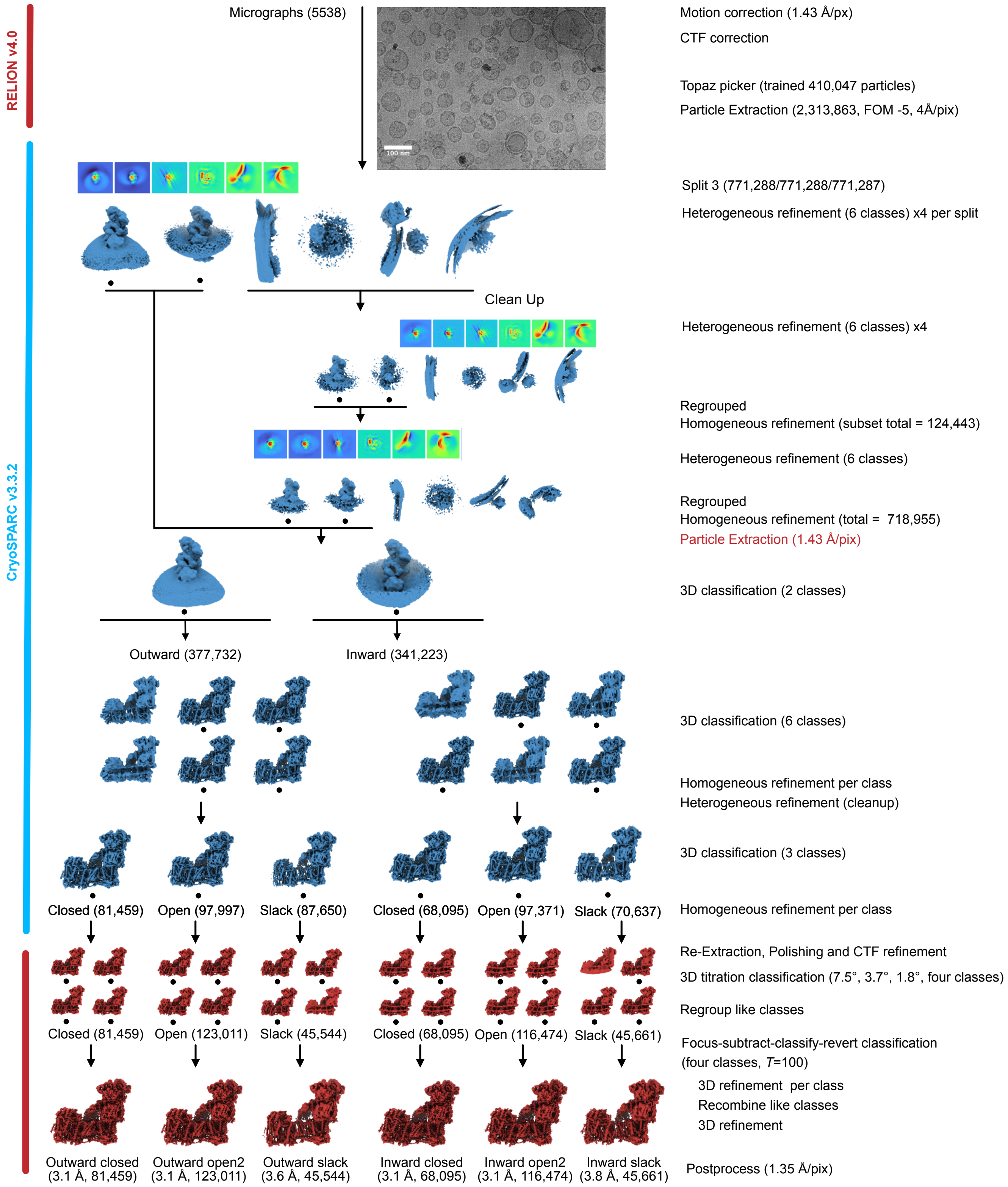


**Fig. S4 | Classification scheme for the DDM-CI-PLs dataset (purified in DDM, as prepared).** An example micrograph and 3D volumes are shown along with cryoEM density heatmaps for volumes in cryoSPARC heterogeneous refinements. Volumes taken forward are indicated with a black dot. The software used to perform each step is indicated by the left-hand colour bar (red text indicates RELION-specific stage exceptions). The focus-subtract-classify-revert classification method is described in more detail in the methods and Fig. S1. The open state was assigned as open2 by comparison with the CI-PLs open1 and open2 states.

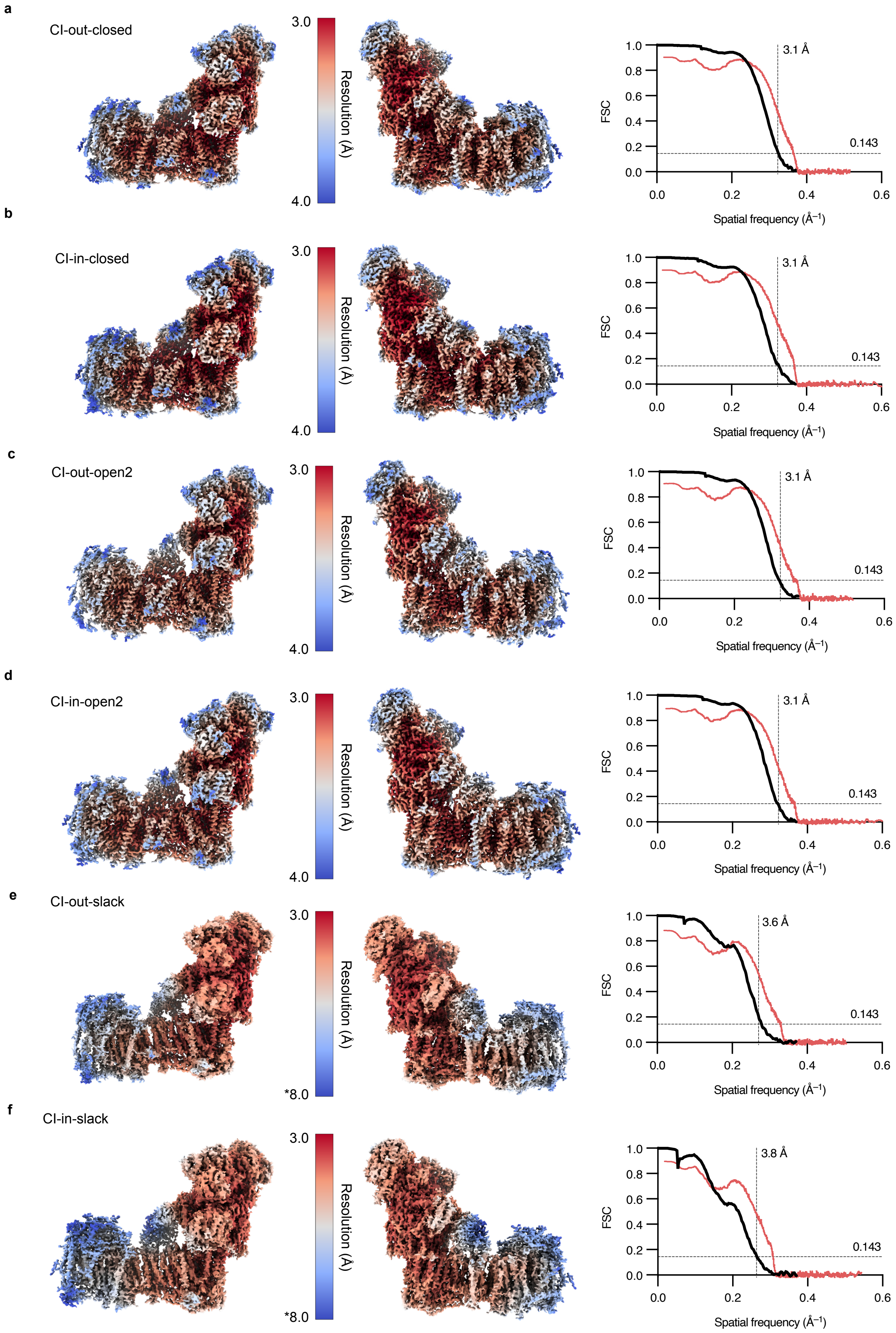


**Fig. S5 | Global and local resolution of the consensus classes of the DDM-CI-PLs dataset (purified in DDM, as prepared).** The final classes **a**) CI-out-closed, **b**) CI-in-closed, **c**) CI-out-open2, **d**) CI-in-open2, **e**) CI-out-slack, and **f**) CI-in-slack were analysed for local resolution distribution using RELION v4.0-beta. Values are plotted according to the colour key. (*) Note for (**e**) CI-out-slack and (**f**) CI-in-slack, the upper value is shifted to 8.0 Å to accommodate the overall lower resolution distribution. Global Fourier Shell Correlation (FSC) scores (cutoff = 0.143) calculated in RELION v4.0-beta are plotted (black) along with model–map FSCs (red), calculated using the validation suite in PHENIX v1.20.1-4487.

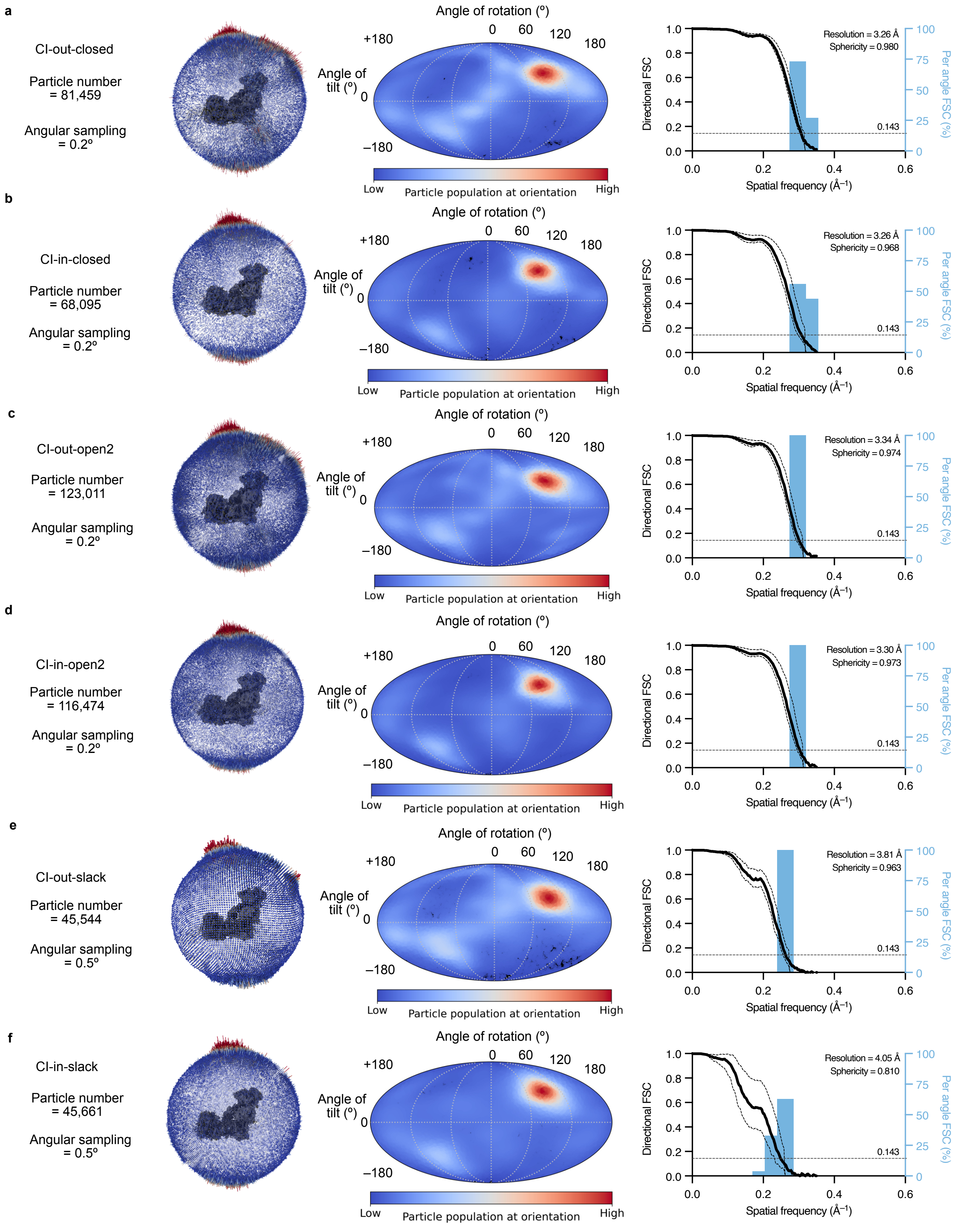


**Fig. S6 | Orientation analyses of the consensus classes of the DDM-CI-PLs dataset (purified in DDM, as prepared).** The final classes **a**) CI-out-closed, **b**) CI-in-closed, **c**) CI-out-open2, **d**) CI-in-open2, **e**) CI-out-slack, and **f**) CI-in-slack were inspected for the orientation distribution of the particles in the final consensus alignment using 3D angular and Mollweide plots. Equivalent data are shown in each case. The orientation distribution in the Mollweide plots is relative to the input reference volume, so the complex I 3D volumes (grey) in each case are shown in the 3D angular distribution spheres for reference. The assessment of uniformity of resolution in each direction (sphericity, where 1 is completely uniform) was performed by dividing the sphere up into 100 evenly sampled cones and calculating directional Fourier-shell correlations (FSCs) plotted as histograms, using the 3DFSC Program Suite v3.0. The 3DFSC global FSC is shown in black (± S.D. of directional FSCs).

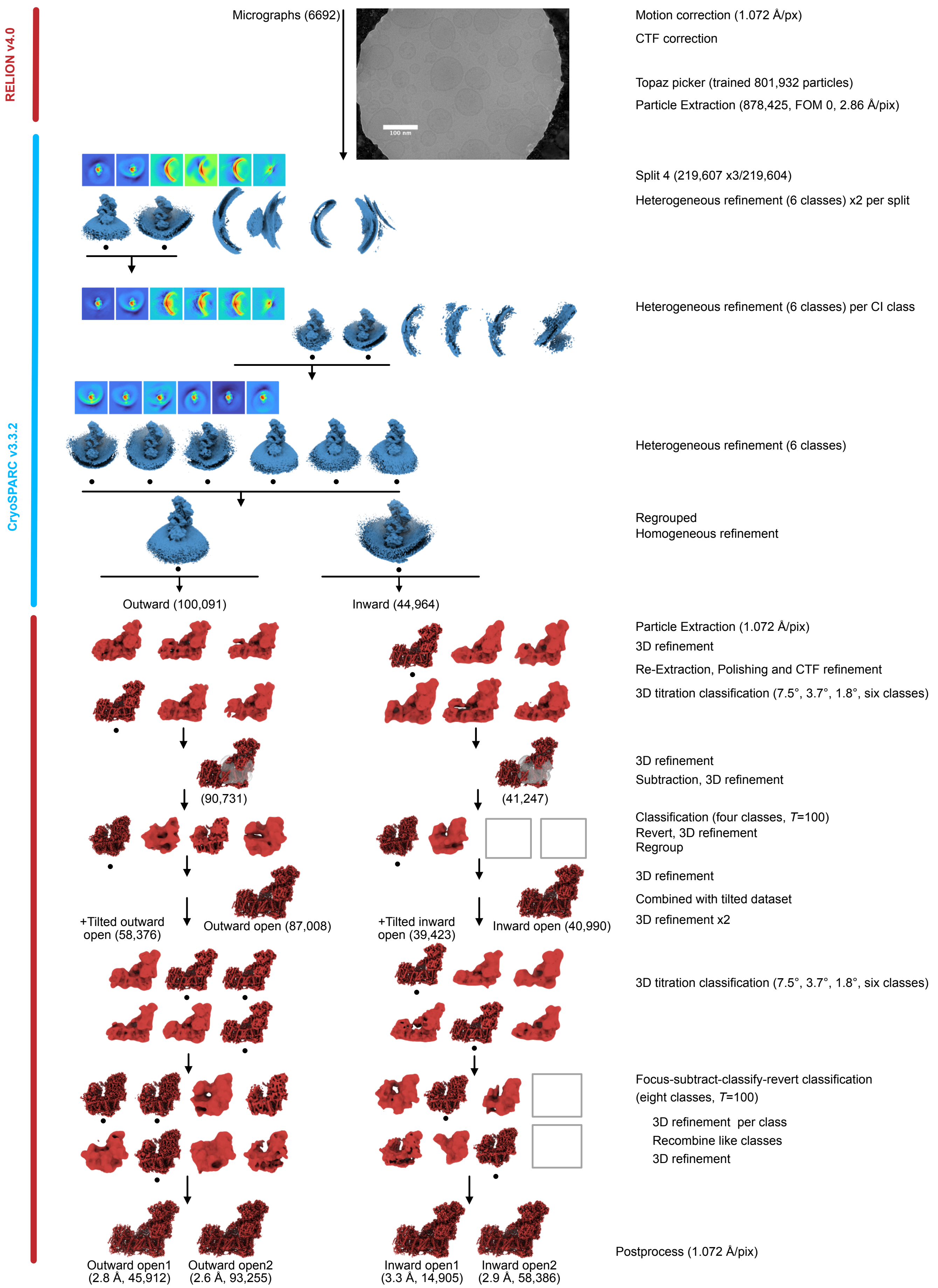


**Fig. S7 | Classification scheme for the non-tilted, deactivated CI-PLs dataset.** An example micrograph and 3D volumes are shown along with cryoEM electron density heatmaps for volumes in cryoSPARC heterogeneous refinements. Volumes taken forward are indicated with a black dot. The software used to perform each step is indicated by the left-hand colour bar. Empty boxes indicate non-populated 3D classes. The focus-subtract-classify-revert classification method is described in more detail in the Methods and Fig. S1. See Fig. S8 for the tilted dataset.

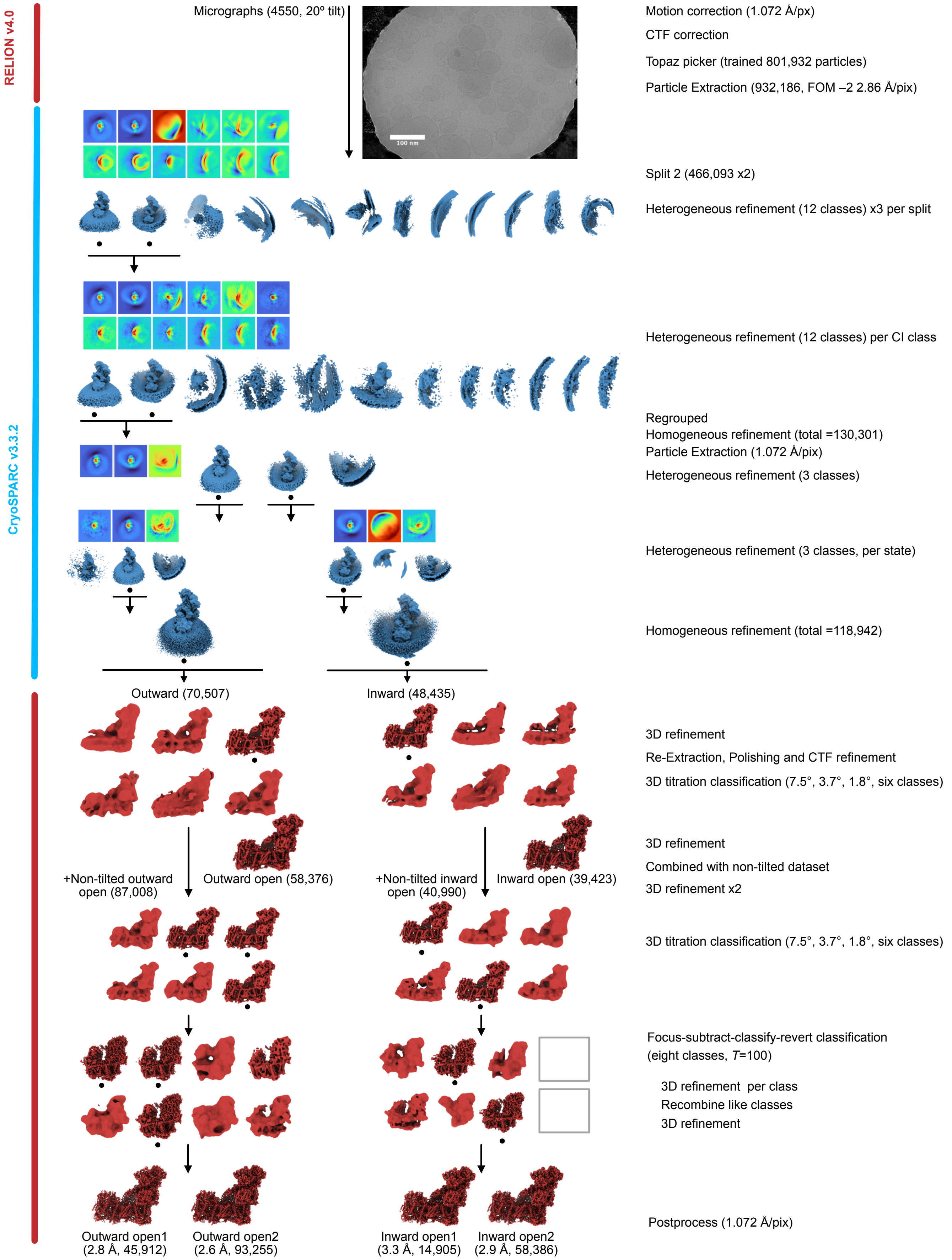


**Fig. S8 | Classification scheme for the tilted, deactivated CI-PLs dataset.** An example micrograph and 3D volumes are shown along with cryoEM electron density heatmaps for volumes in cryoSPARC heterogeneous refinements. Volumes taken forward are indicated with a black dot. The software used to perform each step is indicated by the left-hand colour bar. Empty boxes indicate non-populated 3D classes. The focus-subtract-classify-revert classification method is described in more detail in the Methods and Fig. S1. See Fig. S7 for the non-tilted dataset.

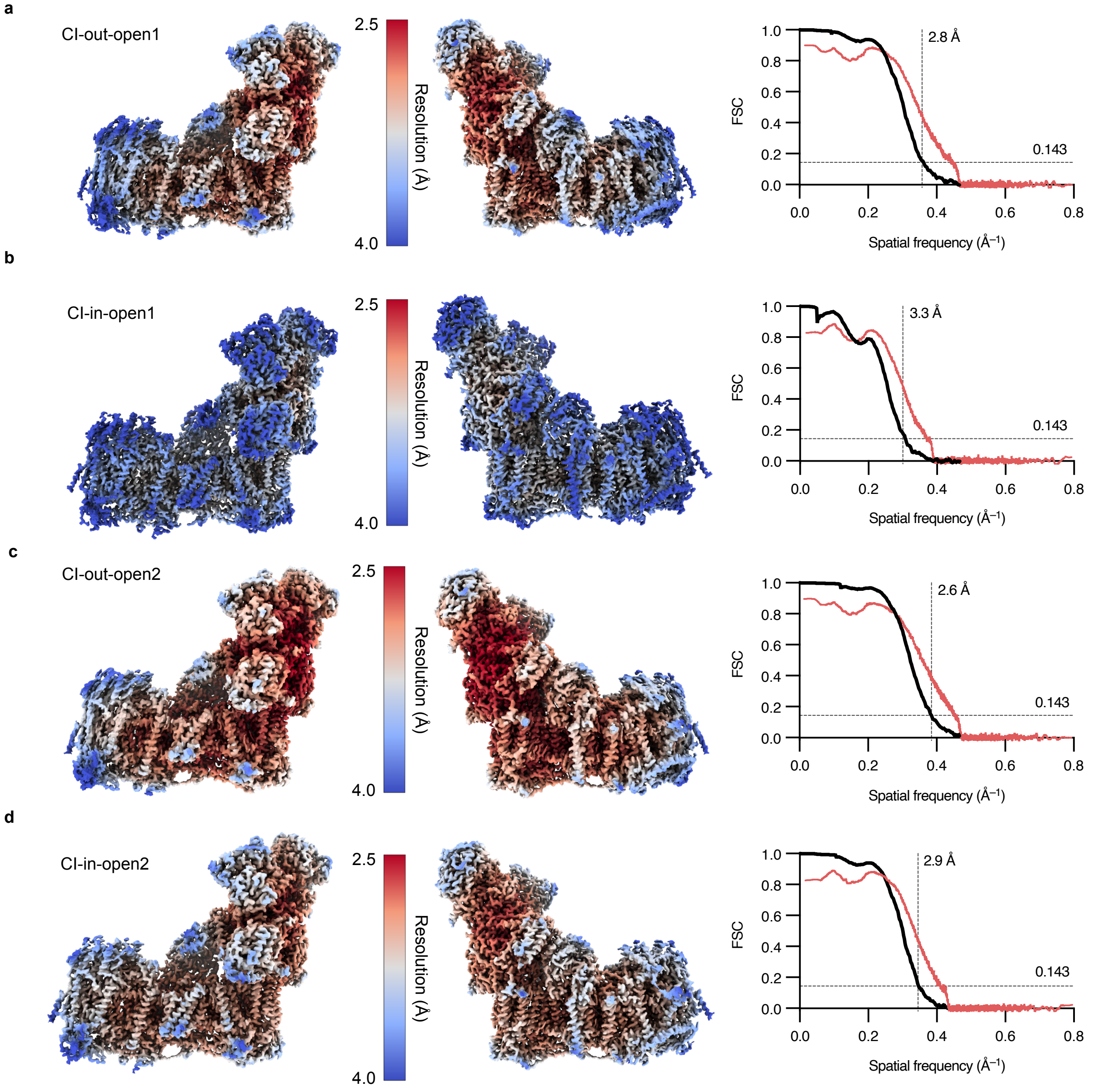


**Fig. S9 | Global and local resolution of the consensus classes of the combined (non-tilted and tilted) deactivated CI-PLs dataset.** The final classes **a**) CI-out-open1, **b**) CI-in-open1, **c**) CI-out-open2, and **d**) CI-in-open2 were analysed for local resolution distribution using RELION v4.0-beta. Values are plotted according to the colour key. Global Fourier Shell Correlation (FSC) scores (cutoff = 0.143) calculated in RELION v4.0-beta are plotted (black) with model–map FSCs (red), calculated using the validation suite in PHENIX v1.20.1-4487.

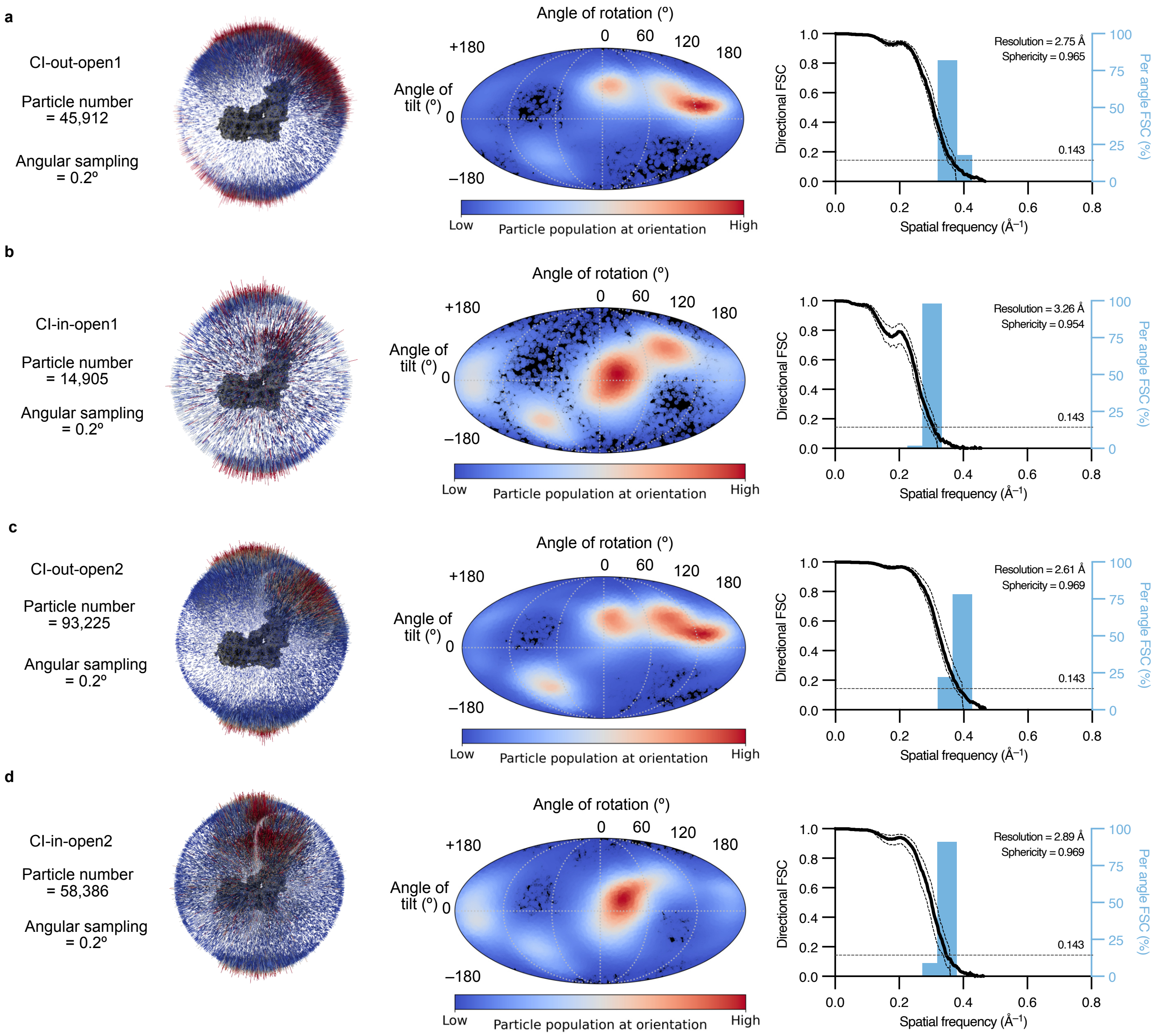


**Fig. S10 | Orientation analyses of the consensus classes of the combined (non-tilted and tilted) deactivated CI-PLs dataset.** The final classes **a**) CI-out-open1, **b**) CI-in-open1, **c**) CI-out-open2, and **d**) CI-in-open2 were inspected for the orientation distribution of the particles in the final consensus alignment using 3D angular and Mollweide plots. Equivalent data are shown in each case. The orientation distribution in the Mollweide plots is relative to the input reference volume, so the complex I 3D volumes (grey) in each case are shown in the 3D angular distribution spheres for reference. The assessment of uniformity of resolution in each direction (sphericity, where 1 is completely uniform) was performed by dividing the sphere up into 100 evenly sampled cones and calculating directional Fourier-shell correlations (FSCs) plotted as histograms, using the 3DFSC Program Suite v3.0. The 3DFSC global FSC curve is shown in black (± S.D. of directional FSCs).

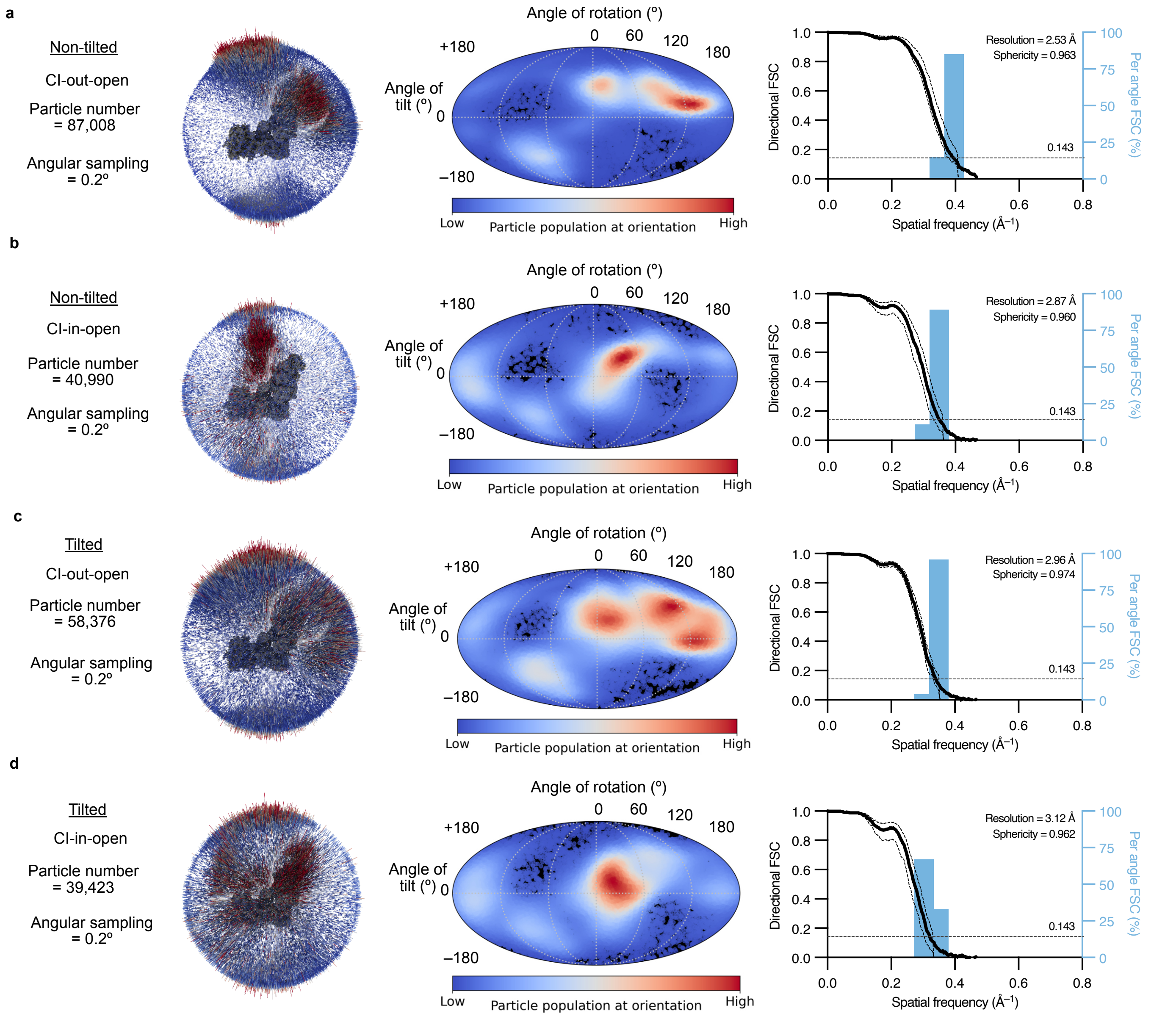


**Fig. S11 | Orientation analyses of the consensus classes of the separately processed non-tilted and tilted deactivated CI-PLs datasets.** The final classes for the non-tilted **a**) CI-out-open, **b**) CI-in-open classes, and the tilted **c**) CI-out-open, and **d**) CI-in-open classes were inspected for the orientation distribution of the particles in the final consensus alignment using 3D angular and Mollweide plots. Equivalent data are shown in each case. Note, at this stage the open1 and open2 states had not been separated and so are presented as a single open class here. The orientation distribution in the Mollweide plots is relative to the input reference volume, so the complex I 3D volumes (grey) in each case are shown in the 3D angular distribution spheres for reference. The assessment of uniformity of resolution in each direction (sphericity, where 1 is completely uniform) was performed by dividing the sphere up into 100 evenly sampled cones and calculating directional Fourier-shell correlations (FSCs) plotted as histograms, using the 3DFSC Program Suite v3.0. The 3DFSC global FSC is shown in black (± S.D. of directional FSCs). The tilted dataset shows increased orientation distribution relative to the non-tilted dataset, however its overall resolution suffers, likely due to the relatively thicker ice and contrast transfer function (CTF) gradient (even with per-particle CTF refinement performed across the micrographs for the tilted grid), as well as increased beam-induced motion. The data presented suggest the level of preferred orientation in the non-tilted dataset does not substantially impact the consensus map, given the number of particles obtained. However, it may negatively impact a reconstruction preformed using a lower number of micrographs/particles.
